# Modeling human B cell development with pluripotent stem cells

**DOI:** 10.64898/2026.05.04.722674

**Authors:** Xiaoning Sun, Jamie J. Kwan, Krishna Kothari, Alexandra F. Nazzari, Astrid Kosters, Colin A. Fields, Bao Q. Thai, Deepta Bhattacharya, Michael Atkins, Kelvin Chan Tung, Xinyuan Zhao, Vladimir T. Manchev, Marion Kennedy, Eliver Ghosn, Gordon Keller

**Author notes:** Current contact. Current address: Faculty of Medicine, Memorial University of Newfoundland, St. John’s, NL. Equal contribution (Jamie J. Kwan, Krishna Kothari, Eliver Ghosn).

## Abstract

The ability to generate functional B cells from human pluripotent stem cells (hPSCs) would open new opportunities to develop novel B cell-based therapies to treat a range of human diseases and disorders. Towards this goal, we established a protocol that promotes the efficient development of B lineage cells from definitive hematopoietic progenitors generated from different hPSC lines. Flow cytometric and multi-omic scRNA-seq analyses revealed that B cell development from hPSCs transitions through the well-established pro-B, pre-B and naïve B cell stages, accurately recapitulating B lymphopoiesis in the human adult bone marrow. Importantly, the naïve B cells generated with this approach could be induced to mature into plasma cells that secrete antibodies and undergo class switching. Analyses of signaling pathways that regulate B lymphopoiesis in these cultures uncovered a potent inhibitory effect of IL-7 on functional IgH rearrangement, resulting in the development of abnormal cells that failed to undergo pre-B cell maturation. Finally, analysis of the different hPSC-derived hematopoietic programs revealed that both definitive and yolk sac progenitors display B cell potential, indicating that there are distinct developmental sources of human B lineage cells. Taken together, these findings demonstrate the efficient generation of B cells from hPSCs and, in doing so, provide a system for further investigating the earliest stages of human B lymphopoiesis and a source of appropriately staged plasma cells for future therapeutic applications.

## Introduction

B lymphoid cells play crucial roles in the innate and adaptive immune systems through their diverse effector functions, including antibody production, complement activation, T cell activation and cytokine secretion.^1^ Beyond these well recognized functions, subsets of B cells have also been shown to display regulatory properties and the capacity to modulate immune responses. Given this diversity, B lineage cells represent potentially important therapeutic targets, offering new opportunities to develop novel treatments for a range of diseases and disorders including protein-deficiencies, infectious, metabolic and autoimmune diseases, cancer and neurological disorders.^2^ One approach that holds great promise is the transplantation of terminally differentiated B cells, the long-lived antibody-secreting cells, or plasma cells, as a cell-based therapy taking advantage of their natural ability to produce and secrete large amounts of protein.^3^ Transplantation of cells engineered to secrete antibodies to specific pathogens could provide long-term immunity to infectious diseases, or those engineered to produce protein drugs, would provide a stable treatment for different monogenic disorders.^4,5^ Towards this goal, several studies have shown that it is possible to transplant engineered peripheral blood-derived plasma cells into recipient mice and that these cells will secrete antibodies for at least several months.^6,7^ While these findings provide proof-of-principle for this strategy, the use of either cord blood- or patient-derived cells has drawbacks that include batch-to-batch variability and limited cell numbers that present challenges for establishing standardized and scalable manufacturing processes. Human pluripotent stem cells (hPSCs) are an attractive alternative to primary cells as they represent a potentially unlimited source of B lineage cells that can be easily genetically modified to produce the desired effector cell of choice as an off-the-shelf, universal cell product. Modeling efficient B-cell development from hPSCs *in vitro,* however, requires an understanding of how this lineage is established in the early embryo and then recapitulating key aspects of this process in the culture dish.

Much of our knowledge of early-life B-cell development comes from studies in the mouse, which show that the B lineage is initially specified in the yolk sac (YS) and then generated in the liver throughout fetal life.^8^ Prior to birth, B lymphopoiesis can be detected primarily in the fetal liver, and after birth, the bone marrow becomes the primary site of B-cell development throughout adult life. Two broad classes of murine B cells, known as B-1 and B-2 B cells, can be distinguished by surface phenotype, anatomical location, function, and developmental origin.^8–10^ B-1 cells are specified early in the yolk sac prior to the emergence of the hematopoietic stem cell (HSC) and colonize the peritoneal and pleural cavities of the animal, where they persist throughout adult life.^11^ Here, a subset of these cells, known as B-1a cells, functions as an innate immune cell population, producing natural antibodies in a T cell-independent manner. B-2 B cells, by contrast, derive from the definitive hematopoietic program and are generated in the late-stage fetal liver and subsequently in the bone marrow throughout adult life. These B cells are found in the bone marrow, spleen, and lymph nodes and represent an essential component of the adaptive immune response. In contrast to the mouse, the onset of human embryonic B lymphopoiesis is not well characterized, largely due to limited accessibility of early embryonic tissue. B-cell progenitors have been identified in the human fetal liver as early as 6 weeks of gestation and were found to express cell-surface phenotypes and gene expression programs distinct from those of their adult bone marrow counterparts.^12,13^ However, the relationship of these fetal progenitors to those found in the adult is unclear, as tracing their early developmental origin and maturation into separate lineages, comparable to mouse B-1 and B-2 B cells, remains a challenge.

B cell development progresses through an ordered series of intermediates beginning with lineage commitment (pre-pro B cells), V-D-J gene rearrangement of the immunoglobulin heavy chain (IgH) locus (pro-B cells), V-J rearrangement of the immunoglobulin light chain (IgL) locus (pre-B cells), and selection of functional B-cell receptor (BCR) repertoire (immature and transitional B cells) prior to maturation (naïve B cells).^14^ Progression through these stages is regulated by waves of specific transcription factors, accessibility of the immunoglobulin loci, expression of the RAG1/2 recombinase genes, cell cycle status, and cytokines such as IL-7. In the mouse, IL-7 has been shown to play a key role in the earliest stages of definitive B-2 cell development, specifically functioning to promote B cell progenitor survival and expansion prior to exit from cell cycle and Ig gene rearrangement.^15^ Mice lacking the IL-7 cytokine fail to develop B-2 cells, but do produce B-1 cells.^16–18^ The role of IL-7 signaling in human B lymphopoiesis, by contrast, is not well understood.^15^ *In vitro* studies have shown that pro/pre-B cell development from both umbilical cord blood and adult marrow CD34^+^ B progenitors is enhanced by the addition of IL-7 to the cultures, suggesting that this pathway may also play a role in human B lymphopoiesis.^19^ However, the observation that humans with IL-7 receptor mutations have normal B-cell numbers suggests that the pathway is not essential for human B lineage development and that other pathways function in its place.^20^

After maturation, naïve B-2 cells remain quiescent until they encounter cognate antigens recognized by their unique B cell receptors (BCRs). Upon antigen recognition, these naïve B cells undergo clonal expansion and differentiation into germinal center B cells for affinity maturation, memory B cells to mount recall responses, or terminal antibody-secreting plasma cells. ^21^ B cell activation and differentiation are driven not only by BCR signals, but also by interactions with CD4^+^ T cells, which provide CD40L and cytokines such as IL-4 and IL-21. Unlike B-2 cells, B-1 cells can be activated without T-cell help and differentiate into plasma cells through activation of innate sensors such as Toll-like receptors.

Advances in our understanding of hPSC differentiation over the past decade have identified key signaling pathways that regulate early embryonic hematopoietic development, enabling the specification of distinct YS and definitive programs.^22–27^ Within the YS branch of the system, we have identified and characterized distinct primitive and multipotent progenitor (MPP) programs, the later likely representing the equivalent of the mouse erythro-myeloid/lymphoid-myeloid (EMP/LMP) programs.^22^ With this resolution, it has been possible to map the origins of different human erythroid, myeloid, and T-cell lineages and demonstrate that their development largely parallels that of the corresponding lineages in the mouse. Detailed analyses of these populations have shown that, as they differentiate, they progress through a progenitor stage known as the hemogenic endothelial cell (HEC), which in turn gives rise to the derivative blood cell types.^23^ While erythroid, myeloid, and T-cell development from hPSCs is well established, the generation of B-lineage cells is not. Several studies have shown that CD19^+^ B-lineage cells can be derived from hPSCs under different culture conditions.^28–32^ However, in most cases, the cells did not progress beyond the pro-B stage of development, and the pathways regulating lineage development were not well defined.

In this study, we developed a protocol to model human B-cell development with hPSC-derived definitive hemogenic endothelial cells as a starting population. We show that these progenitors develop efficiently and transition through the expected stages of B cell development, giving rise to naïve IgD^+^ IgM^+^ B cells that display diverse VDJ gene family usage. Following further culture under appropriate activation conditions, the naïve B cells differentiate into plasma cells that secrete IgM, IgA, and IgG, indicating they can undergo class switching. By manipulating cell signaling pathways in culture, we found that sustained IL-7 signaling inhibited differentiation at the pre-B cell stage, thereby identifying a role for IL-7 signaling in human IgH rearrangement and B cell maturation. Finally, by extending our analyses to YS hematopoiesis, we show that the MPP but not the primitive program also displays B-cell potential, suggesting that there are multiple developmental sources of human B cells.

## Results

### Derivation of B lymphocytes from human cord blood progenitors

To establish optimal conditions for human B lymphopoiesis *in vitro*, we focused our initial studies on cord blood (CB) progenitors, building from the protocol of Laurenti E. et al.^33^ that included a broad range of cytokines (GM-CSF, G-CSF, SCF, TPO, IL-7, IL-6, FLT3L and IL-2) designed to support the generation of lymphoid and myeloid lineage cells following culture on MS-5 stromal cells (Figure S1A). For these optimization experiments, we eliminated factors in a sequential fashion to identify the combination that supported the highest number of B cells and lowest number of NK cells. We focused on these parameters as NK cells, if present in high numbers, can destroy the MS-5 stroma. With this approach, we found that removal of the myeloid factors, GM-CSF and G-CSF (-2G), together with IL-2 resulted in a significant increase in the total number of CD19^+^ cells generated compared to all other factor combinations (Figure 1A, S1B). This combination also generated relatively low number of CD56^+^ NK cells. Removal of FLT3L along with the myeloid factors showed a similar increase in the number of B cells generated. In addition to CD19^+^ B lineage cells, these factor combinations also promoted the development of CD33^+^ myeloid cells to different degrees. B cell frequencies varied more than the total cell number as those in the -2G-IL-2 and -2G-FLT3L cultures were higher than some, but not all other groups (Figure S1C). Taken together, these findings show that we are able to selectively enhance B cell development on MS-5 stroma cells through the manipulation of cytokines and that the combination of TPO, SCF, IL-7, IL-6 and FLT3L (-2G-IL-2) promotes the generation of higher numbers of B cells compared to cultures with most other combinations of factors.

**Figure 1.**
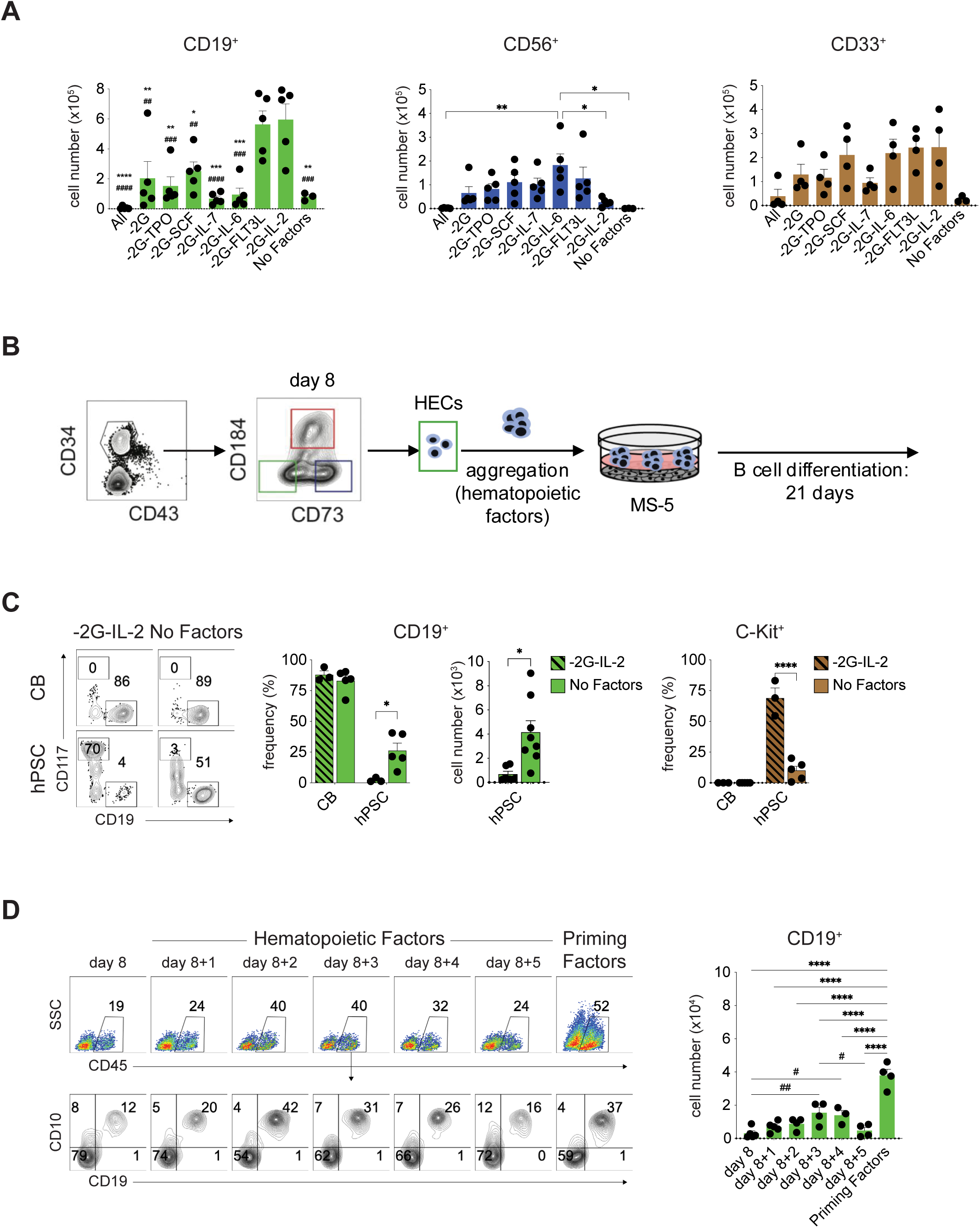
B cell development from cord blood- and hPSC-derived progenitors. (A) Quantification of the number of CD19^+^ B cells, CD56^+^ NK cells and CD33^+^ myeloid cells generated in each of the indicated cytokine combinations. *n* ≥ 3. For evaluation of B cells, (*) represents comparison with -2G-FL3L group and (^#^) represents comparison with -2G-IL-2 group. Values shown are from one well of a 6 well plate seeded with 10^3^ CD34^+^ cord blood cells. Error bars represent SEM. \**P* < 0.05, \*\**P* < 0.01, \*\*\**P* < 0.001, \*\*\*\**P* < 0.0001, ^##^*P* < 0.01, ^###^*P* < 0.001, ^####^*P* < 0.0001 by one way ANOVA analyses with Tukey’s multiple comparisons. (B) Schematic of differentiation protocol for generation of B cells from human PSCs. 10^4^ day 8 CD34^+^CD43^-^CD73^-^ HECs generated from H1 hPSCs were reaggregated for 2 days (unless otherwise indicated) in the broad combination of hematopoietic cytokines (TPO, IL-3, IL-6, IL-11, SCF, EPO, IGF-1, VEGF, bFGF, BMP4) or in the ‘priming factors’ (TPO, IL-3, G-CSF, FLT3L and SCF) in wells of a 96-well low-cluster plate. The resulting aggregates were then co-cultured with MS-5 stroma in the presence of B cell factors TPO, SCF, IL-7, IL-6 and FLT3L (-2G-IL-2) or in the absence of any factors (No Factors) for 21 days. (C) Left: Representative flow cytometric analysis of the frequency of CD19^+^ B cells and CD117^hi^ mast cells generated from either CBs or hPSCs in the presence (-2G-IL-2) and absence (no factors) of factors. Middle and right: Quantification of the frequency CD19^+^ B cells generated from hPSC and CB progenitors, the total number of hPSC-derived CD19^+^ B cells and the frequency of CD117^+^ mast cells generated from both progenitor populations following culture in the presence (-2G-IL-2) and absence (no factors). *n* ≥ 3. Numbers shown are per well of a 6 well plated seeded with either 10^3^ CB cells or the equivalent of 10^4^ day 8 HECs. (D) Left: Representative flow cytometric analysis of B cell development from day 8 HECs reaggregated for different times (0-5 days) in either the broad combination of hematopoietic cytokines (TPO, IL-3, IL-6, IL-11, SCF, EPO, IGF-1, VEGF, bFGF, BMP4) or for 3 days in the priming factors (TPO, IL-3, G-CSF, FLT3L and SCF). The CD10 and CD19 profiles are shown for the gated CD45^+^ cells (upper panel), Right: Quantification of total number of CD19^+^ B cells generated from 10^4^ d8 HECs aggregated for different periods of time or in different media. *n* ≥ 3. Error bars represent SEM. \**P* < 0.05, \*\**P* < 0.01, \*\*\*\**P* < 0.0001, ^#^*P* < 0.05, ^##^*P* < 0.01 by one way ANOVA or two-way ANOVA analyses with Tukey’s multiple comparisons or unpaired *t*-test.

### Generation of B lymphocytes from hPSC-derived hematopoietic progenitors

Using the above optimized protocol, we next investigated the B cell potential of a hPSC-derived CD34^+^ population that we have previously shown displays erythroid, myeloid and T cell potential.^24^ This CD34^+^ population, which develops within 8 days of differentiation, is identified by the lack of expression of CD184 and CD73 that mark arterial and venous endothelial progenitors, respectively, and contains the earliest emerging hematopoietic progenitors, the hemogenic endothelial cells (HECs) (Figure 1B). To assay for B cell potential, day 8 HECs isolated by FACS were aggregated in suspension culture for 2 days under conditions (TPO, IL-3, IL-6, IL-11, SCF, EPO, IGF-1, VEGF, bFGF, BMP4) that we have previously shown promote maturation through a process known as the endothelial-to-hematopoietic transition (EHT).^24^ The progenitors were then cultured on MS-5 stroma in the above optimized conditions (-2G-IL-2) for 21 days. As shown in Figure 1C, the hPSC-derived progenitors did give rise to some CD19^+^ B cells although the frequency was dramatically lower (<5%) than that observed in the CB cultures (>85%). This is likely due, in part, to the high frequency of CD117^+^ mast cells present in these cultures (>70%) that were not detected in the CB-derived populations. As a control, we cultured cells in the absence of any factors and observed quite a different developmental pattern. Under these conditions, the hPSC-derived progenitors generated a significantly higher frequency and total number of CD19^+^ B cells and a lower frequency of mast cells than in the presence of cytokines (Figure 1C). This response is distinct from that of the CB progenitors that produced higher numbers of B cells in the presence of these factors (Figure 1A). Analyses of the CD184^+^CD73^lo^ (arterial; AEC) and CD184^-^CD73^+^ (venous; VEC) fractions within the CD34^+^ population revealed that the B cell progenitors were restricted to the HEC fraction, a pattern similar to that observed for erythroid, myeloid and T cell progenitors (Figure S1D).^24^ Taken together, these findings show it is possible to generate B lineage cells from a hPSC-derived progenitor population and that the optimal conditions for generating these cells are distinct from those for the CB progenitors.

A key step in the efficient generation of hematopoietic progenitors from the day 8 HECs is the aggregation step that promotes the endothelial-to-hematopoietic (EHT) transition. To determine if our conditions were optimal for B progenitor development, we varied the time of aggregation and the cytokine composition of the aggregation media with the goal of enhancing hematopoietic specification and lineage development. As shown in Figure 1D, cells aggregated for 3 or 4 days consistently gave rise to significantly more B cells than the non-aggregated day 8 HECs. Using 3-day aggregates, we next tested different cytokine combinations, specifically focusing on the B-lineage priming cytokines (TPO, IL-3, G-CSF, FLT3L and SCF) identified by Luo et al.^34^ as important for B cell development from cord blood progenitors. This change in cytokines did impact B cell development from the hPSC-derived HECs, as cells aggregated and cultured in the priming factors for 3 days generated significantly more B lineage cells than those aggregated for different periods of time in the combination used in our studies. Given these findings, the day 8 HECs were aggregated for 3 days in the presence of the priming factors prior to MS-5 culture for the following experiment.

### Characterization of the hPSC-derived B lymphocytes

To further characterize hPSC-derived B lineage cells, we next analyzed the temporal patterns of differentiation, monitoring the populations at three different time points (days 7, 14 and 21) for the loss of CD34^+^ expression and the upregulation of expression of CD19 and CD10 and CD19 and CD20, marker combinations that broadly define pro-B and pre-B cells, the earliest stages of B lineage development. Cultures initiated with CB CD34^+^ progenitors were evaluated in parallel. The outcome of these analyses showed similar patterns in both the hPSC and CB cultures that included a decline in the proportion of CD34^+^ progenitors over this two-week period and the emergence of a CD19^+^ population by day 14 (Figure 2A). A portion of these CD19^+^ cells expressed CD10 and/or CD20 at this timepoint. By day 21, the majority of the CD19^+^ cells from both progenitors co-expressed CD10, and this CD19^+^CD10^+^ fraction represented between 36% ± 9.9 (*n* = 3) and 73.9% ± 9.2% (*n* = 7) of the entire CD45^+^ population at this stage. The proportion of the day 21 CD19^+^ population that co-expressed CD20 was considerably lower (average 24.5% ± 12.6 *n* = 3 for hPSC progenitors and average of 43.4% ±14.1, *n* = 4 for CB progenitors). Few IgM^+^ cells were detected at day 21 (not shown), indicating that these populations represent a mixture of CD10^+^CD19^+^ and CD10^+^CD20^+^ pro-/pre- B cells that have not yet progressed to the mature naïve B cell stage of development. CD33^+^ myeloid cells were detected in both the hPSC- and CB-derived populations at all time points analyzed.

**Figure 2.**
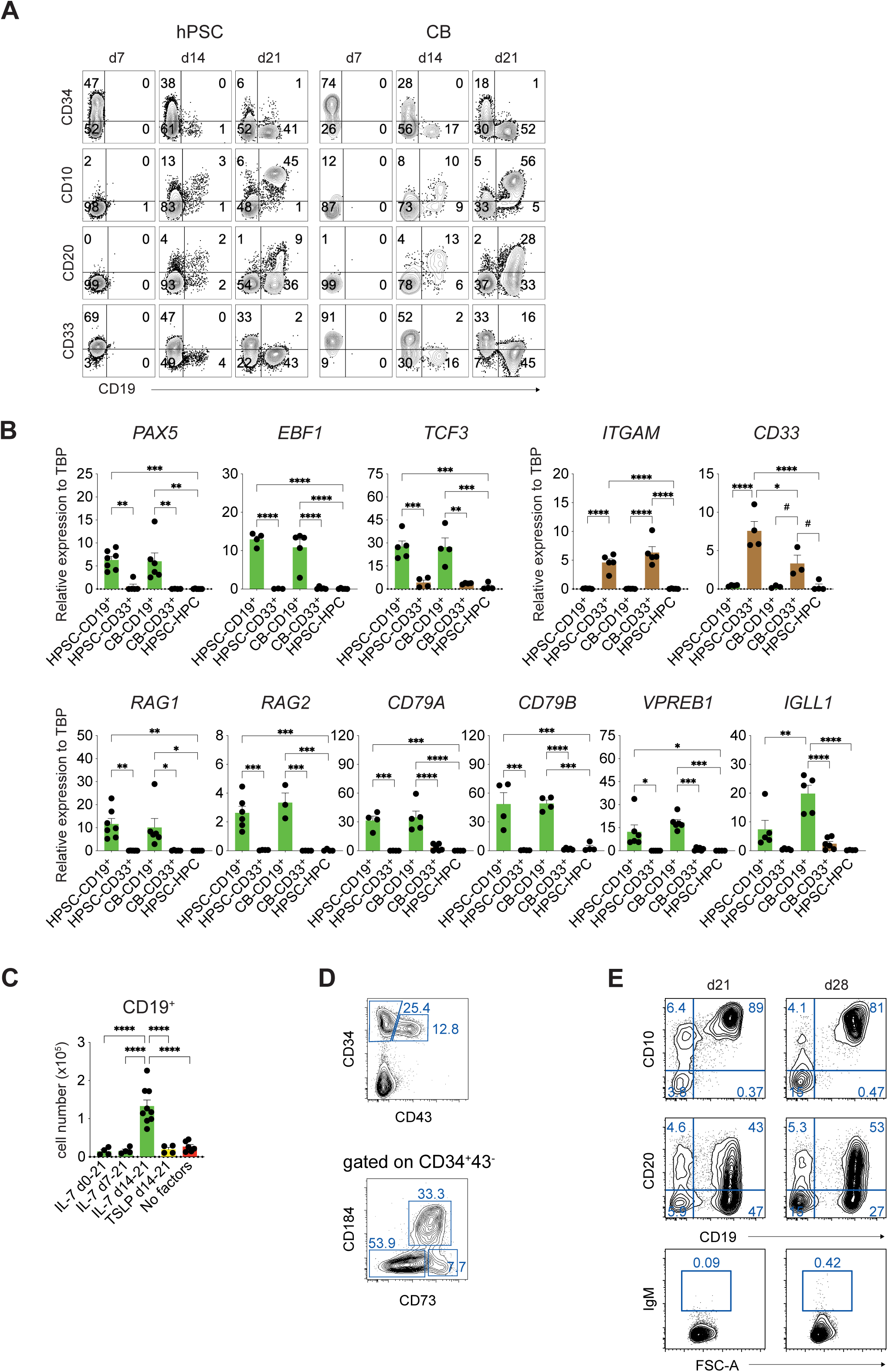
Characterization of hPSC-derived B cell development. (A) Flow cytometric analyses of expression of the indicated markers on hPSC- and CB-derived populations following 7, 14 and 21 days of culture on MS-5. Both hPSC and CB progenitors were cultured in the absence of factors to compare B cell development under the same conditions. hPSC-derived cultures were initiated with 10^4^ day 8 HECs aggregated for 3 days in media containing the priming factors. (B) qPCR analysis of the expression of the indicated genes in FACS isolated day 21 CD19^+^ B cells (green bars) and CD33^+^ myeloid cells (brown bars) generated from hPSC or CB progenitors. D8 HECs or d8+3 aggregates collectively referred to as HPSC-HPCs were used as negative controls. *n* ≥ 3. Error bars represent SEM. \**P* < 0.05, \*\**P* < 0.01, \*\*\**P* < 0.001, \*\*\*\**P* < 0.0001, ^#^*P* < 0.05 by one way ANOVA analyses with Tukey’s multiple comparisons. (C). Quantification of the number of CD19^+^ B cells generated from hPSC-derived progenitors cultured in the indicated conditions. IL-7, TSLP, and No factors treatments are shown by the green, yellow and red bars respectively. Cultures were initiated with 10^4^ day 8 HECs reaggregated for 3 days in priming cytokines *n* ≥ 3. (D) Representative flow cytometric analyses of expression of CD34, CD43, CXCR4 and CD73 on day 13 populations. (E) Representative flow cytometric analyses of expression of the indicated markers on day 13 HEC-derived B cell populations cultured for either 21 or 28 days on MS-5. IL-7 (20 ng/ml) was added on day 14 and maintained throughout culture period.

RT-qPCR analyses showed that the day 21 CD19^+^ cells generated on MS-5 stroma expressed genes associated with the development of the B-cell lineage (Figure 2B). *PAX5*, *EBF1* and *TCF3*, transcription factors known to act in concert to drive B-cell specification and commitment in the mouse^35–37^ were expressed at comparable levels in the hPSC- and CB-derived populations. Both CD19^+^ populations also expressed similar levels of *RAG1* and *RAG2*, genes that encode enzymes required for heavy and light chain V(D)J gene rearrangements^38^, *CD79A* (Igα) and *CD79B* (Igβ), genes that encode proteins associated with the B-cell antigen receptor (BCR) and *VPREB1* (*CD179A*), a component of the pre-BCR surrogate light chain^39^. The CB cells expressed higher levels of *IGLL1* (λ5) (*CD179B*) the second component of the pre-BCR.^40^ As expected, the CD33^+^ myeloid cells did not express appreciable levels of these B lineage genes but did express the myeloid genes *ITGAM* (CD11b) and *CD33*. Collectively, these expression patterns support the findings from the flow cytometric analyses that the day 21 populations generated from hPSC-derived and CB progenitors represent the pro- and pre-B cell stages of development.

### Optimization of B-cell development from hPSCs

Although our findings show that it is possible to generate B lineage cells from hPSCs in the absence of added factors, the efficiency of differentiation under these conditions was low, with typical yields of between 1x10^4^ and 5x10^4^ CD19^+^ cells per 10^4^ input HECs. To improve B-cell output in this model, we investigated two different aspects of the culture system. The first was to determine if the staged manipulation of specific signaling pathways would have any effect on the number of CD19^+^ cells generated. Here, we focused on IL-7 signaling. The second was to investigate different stages of EB differentiation, with the aim of identifying a more enriched B-cell progenitor population.

For analyses of the effect of IL-7, the cytokine was added to the cultures at different time intervals to identify potential stage-specific responsiveness. The intervals included addition either throughout the 21 days culture period (0-21), from day 7-21 or from day 14-21 of culture (Figure S2A). Addition of IL-7 throughout the culture (0-21) or from day 7 onward led to a significant reduction in the frequency of CD19^+^ cells and to an increase in both the frequency and total number of CD56^+^ NK cells compared to cultures without IL-7 (Figure 2C, S2B). In contrast, addition of IL-7 at day 14 led to a significant increase (approximately 10-fold) in both the frequency and total number of CD19^+^ cells generated compared to addition at the other times or to the no factor control (Figure 2C). The late addition of IL-7 also reduced the frequency of CD56^+^ NK cells generated compared to the populations treated at day 7 or for the entire culture period (Figure S2B). Addition of thymic stromal lymphopoietin (TSLP), which promotes signaling through IL7Rα-TSLPR heteromers, had no impact on the frequency or total number of CD19^+^ cells (Figure 2C, S2B).

Analyses of the different stages of EB differentiation revealed that the CD34^+^CD43^-^CD184^-^CD73^-^ HEC fraction persisted over time and by day 13 represented from 25% to greater than 50% [40 +/- 4.5% (SEM), N=7] of the entire CD34^+^CD43^-^ population (Figure 2D). At this timepoint the CD34^+^CD43^-^ fraction made up from 10% to 25% [19 +/- 3.0% (SEM), N=7] of the whole EB population. As the day 13 HECs aggregated poorly, they were cultured directly on the MS-5 stroma in the presence of the priming factors (Figure S2C). IL-7 was added to the cultures between day 14 and 21. This day 13 progenitor population did generate CD19^+^CD10^+^ and CD19^+^CD20^+^ pro/pre-B cells and under these conditions and did so at much higher efficiency than the day 8 progenitors as we were able to routinely generate populations, consisting of up to 80% CD19^+^CD10^+^ cells by day 21 of culture (Figure 2E). The fact that the reaggregation step was not necessary indicates that EHT likely occurs in the EBs prior to the isolation of the CD34^+^CD43^-^CD184^-^CD73^-^fraction. Given these efficiencies, we used the day 13 HEC population for all subsequent experiments.

To promote further maturation of the pre-B cells, we maintained the population in culture for an additional week in the presence of IL-7 (Figure 2E). These conditions supported the CD19^+^CD10^+^ and CD19^+^CD20^+^ populations but failed to promote any further maturation as few IgM^+^IgD^+^ cells were detected. The relatively high frequency of CD19^+^CD20^+^ cells in populations with virtually no IgM^+^ cells was unexpected and suggested that the B-lineage cells may not be following the normal pattern of development under these conditions.

### Molecular analyses of IL-7 treated hPSC-derived B cells

To investigate the potential developmental block observed in the presence of IL-7, we performed multi-omic scRNA-seq analyses on the hPSC-derived day 21 CD19^+^ population cultured in the presence of IL-7 (20 ng/ml) from day 14 to 21. This method enables analyses of global transcriptomics, Ig heavy and light chain V(D)J rearrangements and cell-surface marker expression at the single-cell level, as previously described(Figure 3A).^41^ A primary human adult bone marrow B-cell sample was analyzed as a control for normal B-cell development. With this approach, we identified six populations of CD19^+^ B cells that correspond to the progressive stages of B cell development, including subpopulations of pro-B cells, pre-B cells, immature B cells and mature B cells (Figure 3B). Integrating the bone marrow-derived and hPSC-derived B cell samples enabled stage-specific comparisons. Using the bone marrow as a reference, the hPSC-derived progenitors appear to follow the expected trajectory of development through the pro- and pre-B cell stages (clusters 1-4) depicted in the UMAPs (Figure 3C).

**Figure 3.**
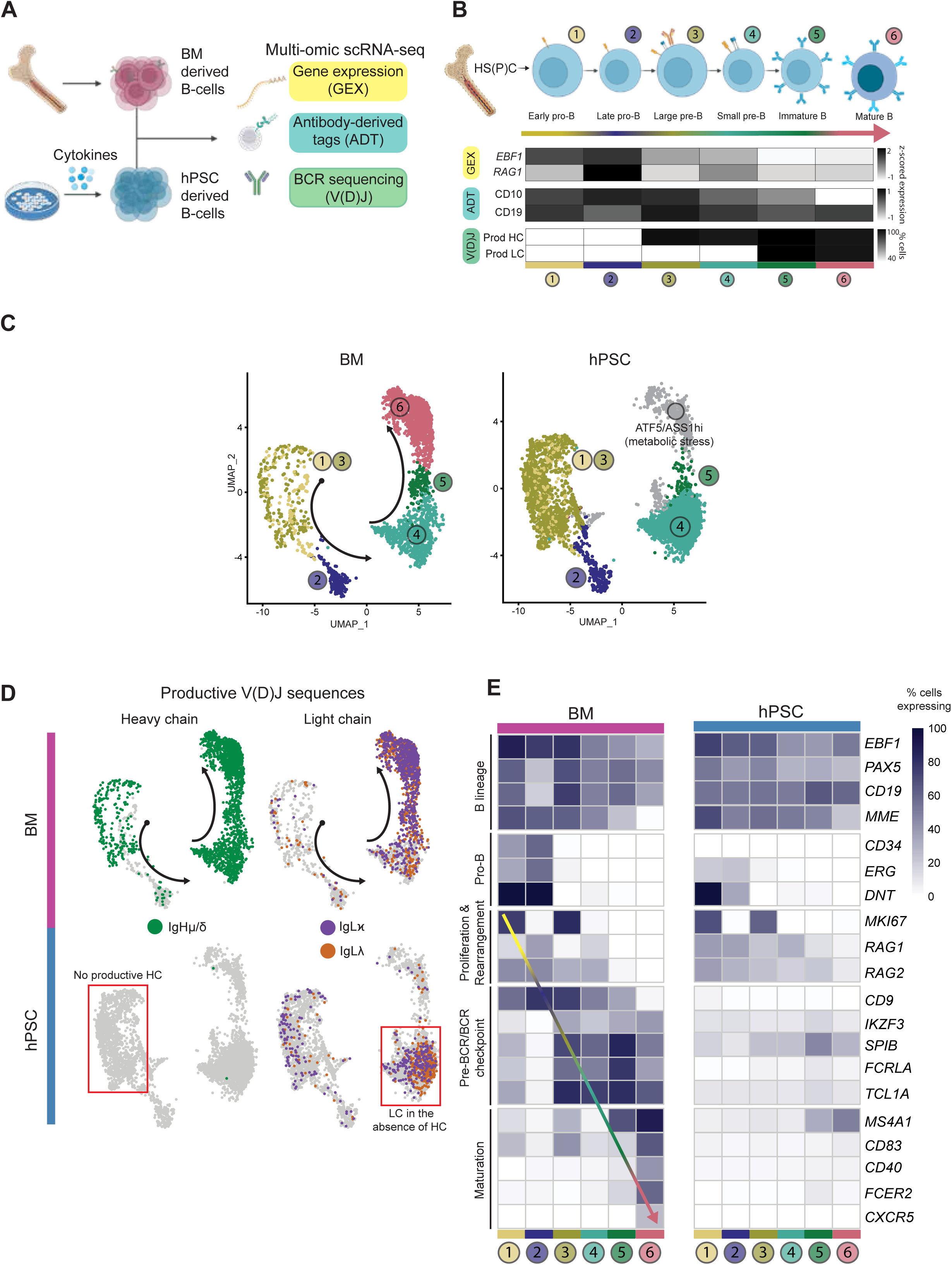
Multi-omic scRNA-seq analyses of adult bone marrow-derived and IL-7 treated hPSC-derived B lineage cells. (A) Experimental design to generate multi-omics single-cell sequencing from hPSC-derived B cells and from human primary B cells isolated from adult bone marrow donors (B) Characterization of the adult bone marrow B-cell developmental pathway based on total gene expression (GEX), cell-surface markers using antibody-derived tags (ADT), and B-cell receptor Heavy and Light chain sequences (V(D)J). Prod HC: productive Heavy Chain; Prod LC: productive Light Chain. (C) Harmony-integrated UMAP (Uniform Manifold Approximation and Projection) of adult bone marrow B and hPSC-derived B lymphopoiesis. The bone marrow data include CD79A^+^ cells (BM, n=2,504 cells) isolated from two adult donors and the hPSC data are from CD19^+^ cells isolated from the MS-5 stroma (magnetic bead sorted) at day 21 of culture. IL-7 was added between days 14 and 21 of culture (IL-7 treated hPSC, n=4,974 cells). Clusters are numbered according to the developmental stages shown in ‘B’. Arrows indicate the direction of B-cell development as determined by the 3-OMIC characterization. (D) UMAP depicting cells expressing productive immunoglobulin heavy chain (IgH; green) and light chain (IgL), kappa (purple) or lambda (orange) (IgL𝜅 or IgL𝜆), in the human bone marrow (BM) and hPSC (IL-7 treated hPSC) B cell populations. hPSC-derived B cells fail to generate productive Heavy Chain (HC) in the Late pro-B and Large pre-B stages (clusters 2 & 3), despite rearranging their Light Chain (LC). (E) Heatmap of the percentage of cells expressing selected genes (GEX) across the B-cell developmental stages (Clusters 1 to 6). Genes are grouped to highlight stages in B-cell development. A direct comparison between human BM and hPSC-derived B cells, as shown in the heatmap, reveals pre-BCR/BCR checkpoint genes that are absent or dysregulated in hPSC-derived cells.

Notably, the hPSC-derived population lacked robust populations beyond the small pre-B stage with few cells at the immature and mature B-cell stages (clusters 5-6), a pattern consistent with the lack of IgM^+^ cells in these cultures detected by flow cytometry (Fig 2E). Rather than displaying a mature phenotype, the hPSC-derived population in cluster 6 contained cells that expressed genes associated with metabolic stress including *ATF5* and *ASS1*, indicative of *abnormal* development. BCR sequencing analyses provided further insights into the aberrant development and the differentiation block. The BM-derived B cells display the expected progression of Ig V(D)J rearrangement such that rearrangement events in pro-B cells result in productive heavy chain transcripts and that in small pre-B cells results in productive light chain (lambda or kappa) transcripts. Thus, immature and mature B cells express productive heavy and light chain rearrangements as well as gene expression patterns consistent with maturation, including expression of the surface markers *CD20* (MS4A1), *CD83* and *CD40* among others (Figures 3D, 3E). In contrast, none of the cells in the hPSC-derived population contained productive heavy chain transcripts. Surprisingly, some of these cells did show productive light chain rearrangement in the absence of heavy chains, a pattern not observed in normal B cell development. Given the lack of heavy chains, a failure to form a functional pre-BCR complex at the large pre-B stage is expected (cluster 3). Signaling through the pre-BCR induces proliferation, which must be halted before rearrangement continues, a mechanism that preserves genomic stability.^42^ Genes involved in the pre-BCR checkpoint, including *IKZF3*, *SPIB*, and *TCL1A*^43–45^ showed reduced levels of expression in the hPSC-derived B cells (Figure 3E, S3) and most genes associated with maturation were absent (Figure 3E), consistent with the observed block in differentiation. The fact that the pre-BCR program is not upregulated suggests that the MS-5 environment with addition of high concentrations of exogenous IL-7 inhibits heavy chain rearrangement while maintaining the cells in an abnormal proliferative stressed state, preventing the exit from the cell cycle that is required for immunoglobulin gene rearrangement and progression along the normal differentiation path.^15^

### Regulation of B cell development in MS-5 cultures

To determine if IL-7 signaling does negatively impact B-cell maturation in the MS-5 cultures, we next cultured cells in different concentrations of IL-7 or in the absence of factor from days 14 to 21. Analysis of the day 21 populations showed that high concentrations increased the frequency of CD19^+^CD20^+^ cells (Figure S4A). At this point, IL-7 was removed from all cultures, and the cells maintained on MS-5 in the absence of additional factors for 28 or 35 days at which time the populations were harvested and analyzed for the presence of IgM^+^IgD^+^ cells. As shown in Figure 4A, we did indeed observe a strong inhibitory effect of IL-7 on pre-B cell maturation, as in the absence of factor or in the presence of low concentrations, the cells matured and gave rise to IgM^+^IgD^+^ cells by day 28 of differentiation. The development of IgM^+^IgD^+^ cells was completely inhibited for the duration of the culture (to day 35) in populations treated with high concentrations of IL-7 from days 14-21 (Figure 4B, S4B). The observation that the inhibitory effect was observed a full 2 weeks following removal of IL-7 strongly suggests that signaling through this pathway during the pre-B stage of development initiates changes that lead to a permanent block in maturation. The effect of IL-7 appeared to be specific to the maturation of the pre-B cells, as the total number of CD19^+^CD10^+^ cells was not dramatically impacted by the presence of different concentrations of the cytokine (Figure 4C, S4B).

**Figure 4.**
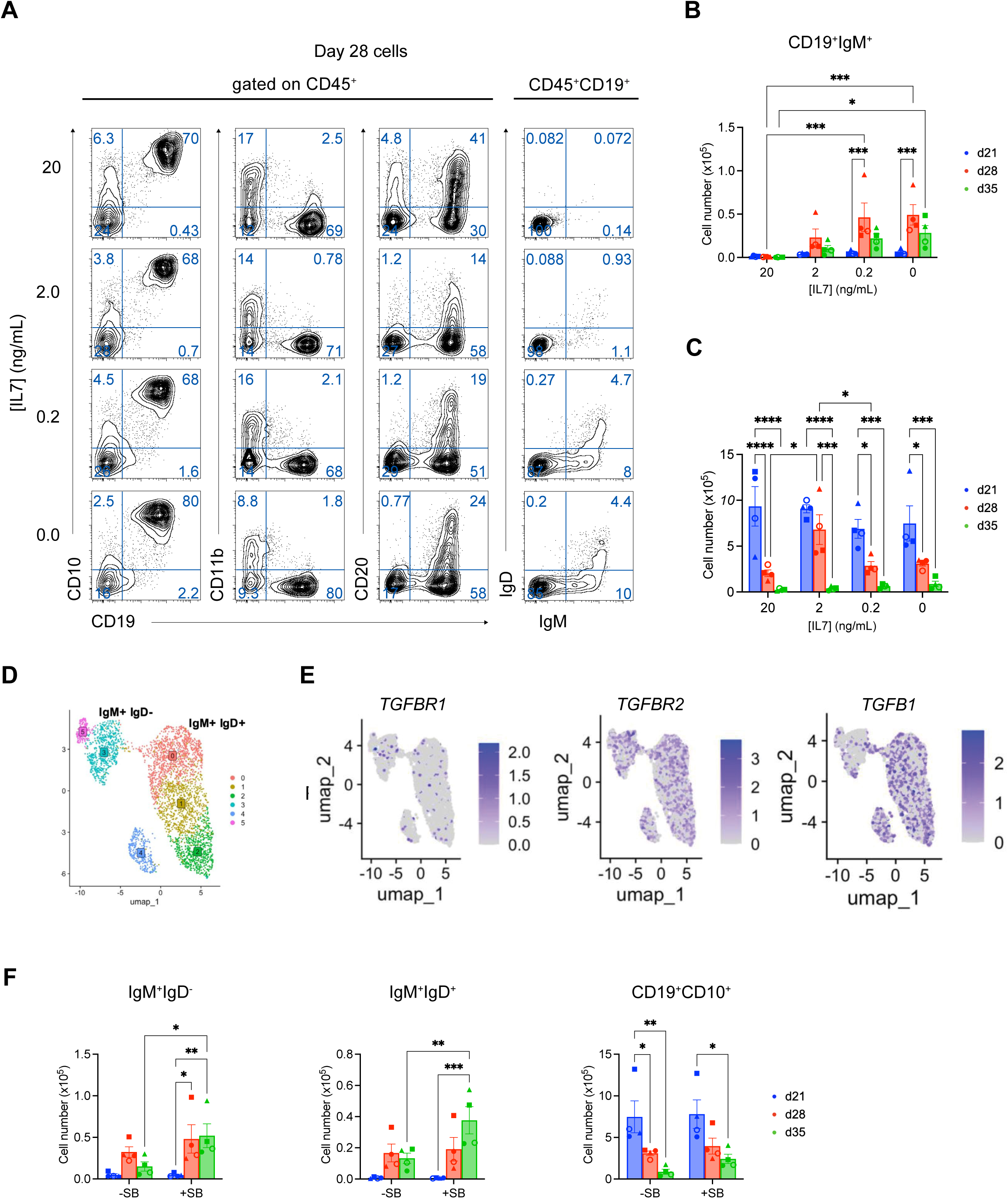
Regulation of B cell development in the MS-5 cultures. (A) Representative flow cytometric analyses of expression of indicated surface markers in hPSC-derived populations cultured for 28 days in the absence or presence of decreasing concentrations of IL-7. IL7 was included in the cultures between days 14 and 21. (B, C) Quantification of the total number of CD19^+^IgM^+^ cells (B) and CD19^+^CD10^+^ cells (C) generated at days 21, 28 and 35 of co-culture. Values shown are per well of a 6-well plate seeded with 4500 day 13 CD34^+^CD184^-^CD73^-^ HECs. Error bars represent SEM. \**P* < 0.05, \*\*\**P* < 0.001, \*\*\*\**P* < 0.0001 by two-way ANOVA analyses with Tukey’s multiple comparisons. Different symbols represent B cells generated from three independent HEC populations, the same symbols (open and closed) indicate B cells generated from the same HEC population (D) UMAP clustering of human CD19^+^ bone marrow cells. Surface expression of IgM and IgD (from TotalSeq antibodies) was used to annotate immature (IgM^+^IgD^-^) and mature (IgM^+^IgD^+^) B cell populations. (E) Feature plots showing expression of TGF-β superfamily receptors grouped by type. (F) Quantification of the number of IgM^+^IgD^-^, IgM^+^IgD^+^ and CD19^+^CD10^+^ cells generated at days 28 and 35 in hPSC-derived populations cultured in the absence or presence of SB. SB was added at day 21 and maintained throughout the entire culture period. Numbers shown are per well of a 6-well plated seeded with 4500 day 13 CD34^+^CD184^-^CD73^-^ HECs. Different symbols represent B cells generated from three independent HEC populations, the same symbols (open and closed) indicate B cells generated from the same HEC population. Error bars represent SEM. \**P* < 0.05, \*\**P* < 0.01, \*\*\**P* < 0.001, by two-way ANOVA analyses with Tukey’s multiple comparisons. N=4

The above findings showed that the hPSC-derived pre-B cells can develop into immature IgM^+^IgD^-^ cells in the presence of fetal calf serum on MS-5 stromal cells in the absence of any other exogenous factors, though the generation of more mature IgM^+^IgD^+^ B cells was modest. To identify other potential regulators of B-cell proliferation and maturation that could improve hPSC-derived B-cell numbers, we analyzed CITE-seq (Cellular Indexing of Transcriptomes and Epitopes by sequencing) data from adult human bone marrow CD19^+^ B cells, focusing on expression of receptors for cytokines known to affect B-cell development and/or proliferation. This approach combines single-cell RNA sequencing with simultaneous cell-surface protein quantification using antibody-derived tags (ADTs) and enables integration of transcriptomic data with direct protein-level readouts of IgM and IgD. Clustering analyses of the data identified six distinct B cell clusters, and IgM and IgD ADT levels revealed that the transition from IgM^+^IgD^-^ to IgM^+^IgD^+^ cells occurred between cluster 3 and 0 (Figures 4D S4C). Signaling pathway analyses showed that these BM-derived B cells express components of the TGFβ signaling pathway, including *TGFBR1, TGFBR2* and *TGFβ1* (Figure 4E). These patterns were of particular interest, as TGFβ signaling has been shown to antagonize human B-cell proliferation *in vitro.*^46^ Analyses of our day 21 hPSC-derived pre-B cell populations showed that they also expressed the same TGFβ receptor and TGFβ1 genes (Figure S4D). These patterns suggest that TGFβ signaling may be playing a similar inhibitory role in the hPSC-derived cultures. To test this, we added either SB431542 (TGFβ inhibitor) or LDN193189 (BMP inhibitor) as a control to the cultures from day 21 to day 35 (Figure S2C). The addition of LDN had no discernible effect on B-cell maturation (not shown). The addition of SB, by contrast, led to a significant increase in the total number of both IgM^+^IgD^-^ (3.5-fold) and IgM^+^IgD^+^ (2.8-fold) cells detected in the populations at day 35 of culture (Figure 4F). The frequency of IgM^+^IgD^-^ cells was also modestly higher in the population cultured in the presence of SB (Figure S4E). The total number of cells and the total number of CD10^+^CD19^+^ cells generated were not impacted by SB, indicating that TGFβ signaling specifically limits the generation of naïve B cells in the MS-5 cultures.

### Temporal development of hPSC-derived B cells

Using these conditions, we next carried out an extended kinetic study to monitor the development and maturation of the hPSC-derived B-cell population over time. The hPSC-derived B-cell lineage developed along the expected trajectory with CD20^hi^ and IgM^+^ cells emerging by day 28 (Figure 5A, B). IgM^+^IgD^+^ cells were readily detected by day 35 of culture and persisted to day 40. The development of kappa and lambda light chains and CD40-positive cells paralleled that of the IgM^+^ cells. During this time course, the frequency and number of total IgM^+^ and IgM^+^IgD^+^ cells increased significantly between days 21 and 35 with some modest, but not significant changes observed between days 35 and 40 in some experiments. These increases in B cells were associated with a decrease in the total number of CD19^+^CD10^+^ cells detected at the later time points. With this protocol, it is possible to generate an average of 14 ± 4 (SEM; n=5), 19 ± 4 (SEM, N=5) and 8 ± 1 (SEM, N=5) IgM^+^ cells per input HEC at days 28, 35, and 40 of culture, respectively.

**Figure 5.**
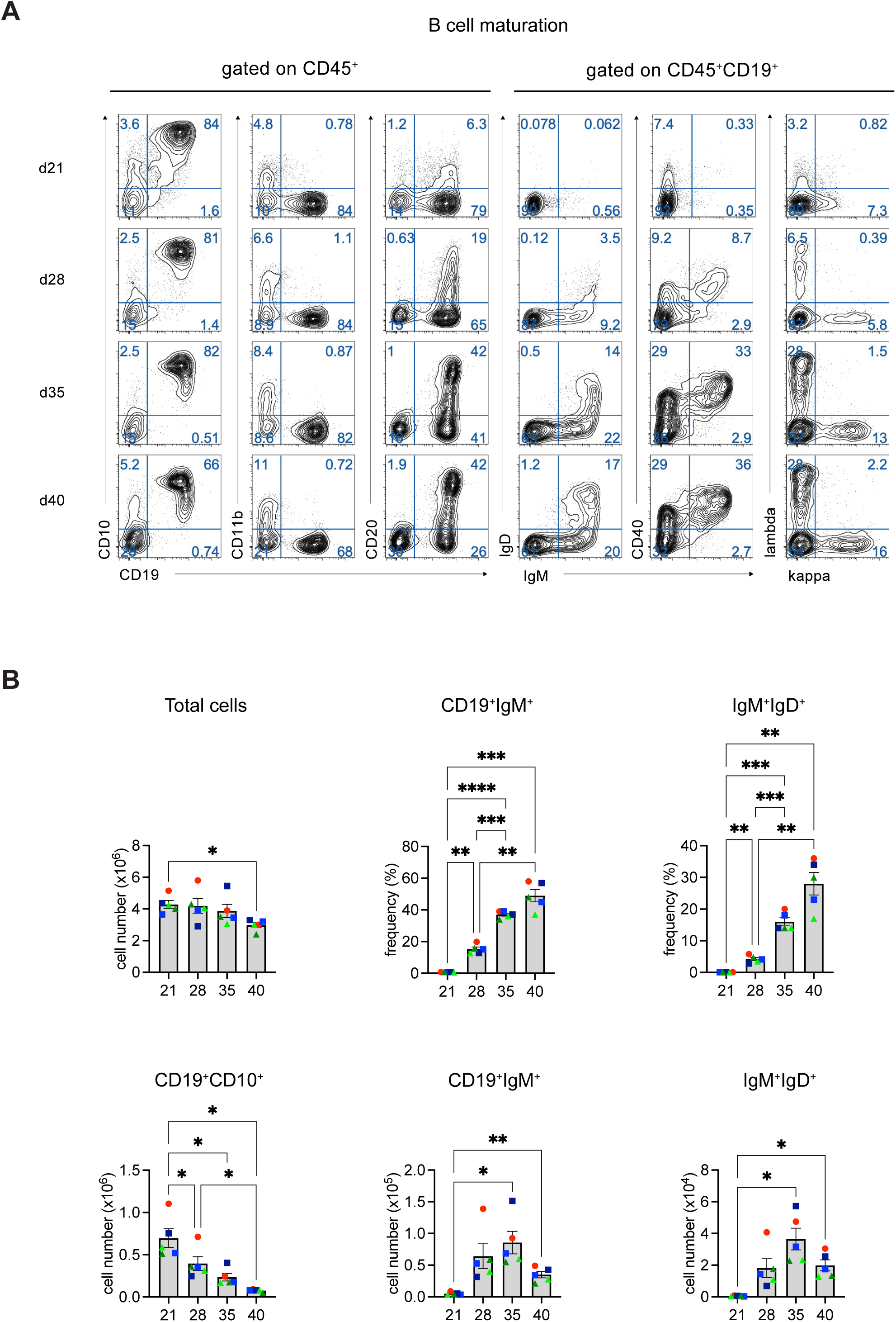
Temporal analyses of hPSC-derived B cell development. (A) Representative flow cytometric analyses of expression of indicated markers on hPSC-derived populations cultured for 21, 28, 35 and 40 days on MS-5 stromal cells. (B) Quantification of the number of total cells, CD19^+^CD10^+^ cells, CD19^+^IgM^+^ cells and IgM^+^IgD^+^ cells and of the frequencies of the CD19^+^IgM^+^ and IgM^+^IgD^+^ populations on the different days. Numbers shown are per well of a 6-well plated seeded with 4500 day 13 CD34^+^CD184^-^CD73^-^ HECs. Different symbols represent B cells generated from three independent HEC populations, the same symbols with different colors indicate B cells generated from the same HEC population. Error bars represent SEM. \**P* < 0.05, \*\**P* < 0.01, \*\*\**P* < 0.001, by two-way ANOVA analyses with Tukey’s multiple comparisons. N=5

To test the B-cell potential across different hPSC lines, we next assayed day 14 HECs generated from either CHOP18^47^ or RM Tomato (RM TOM)^26^ iPSCs (Figure S5A-D). HECs from both iPSC lines generated B lineage cells with a development progression similar to that observed with the H1-derived progenitors. IgM^+^ cells co-expressing kappa^+^ or lambda^+^ light chains emerged by day 28 of differentiation and by day 35 IgM^+^IgD^+^ cells were detected in both populations. The efficiency of B-cell development from these iPSC-derived HEC populations differed from that of H1. With the CHOP line, we generated an average of 2 ± 1 (SEM; N=5) and 1 ± 0.5 (SEM; N=5) IgM^+^ cells per input HEC at days 28 and 35 of culture, respectively. The number was considerably higher with the RM TOM-derived progenitors, as they give rise to 31 ± 11 (SEM; N=6) IgM^+^ cells at day 28 of culture and 65 ± 16 (SEM; N=6) IgM^+^ cells at day 35.

### Characterization and maturation of hPSC-derived naïve B cells

To further characterize the hPSC-derived B cells generated in the absence of exogenous IL-7, we evaluated the day 35 population using the multi-omic scRNA-seq approach. The outcome of these analyses revealed a different picture from the day 21 IL-7-treated population. These cells follow the expected B cell developmental trajectory and successfully rearranged heavy and light chains, as evidenced by the productive Ig transcripts (Figure 6A, 6B). As predicted from the flow cytometric analyses, the day 35 hPSC-derived population contained mature B cells (cluster 6) with productive heavy- and light-chain V(D)J gene rearrangements (Figure 6A, 6B). This population also expressed elevated levels of the transcription factor (*IKZF3*, *SPIB,* and *TCL1A*) and maturation marker (*MS4A1*, *CD83*, and *CD40*) genes (Figure 6C, S6A,C), reflecting the phenotype of canonical B-cell development in the human BM (Figure 3E, 6C). In addition to the mature B cells, the day 35 population also contained a small number of cells (cluster 7) that expressed genes associated with plasma cell differentiation, including *SLAMF7* and IgG class-switching (*IGHG*), demonstrating that the cultures support both differentiation of the B cell lineage and maturation of the naïve B cell to the plasma cell stage (Figure 6D). The Ig V(D)J rearrangement analyses (Figure 6E) showed that, under the given culture conditions, sequential heavy- and light-chain gene rearrangement is permissible, leading to a diverse VDJ-gene family usage similar to a canonical naïve B-cell phenotype. The diverse set of *IGHV* (Figure 6E), *IGHD*, and *IGHJ* (Figure S6B) genes shows that these naïve B cells arise from *de novo* development and maturation in the culture system rather than clonal expansion of a limited number of pre-B cells. Taken together, the findings from these molecular analyses are in line with the flow cytometry patterns and show that it is possible to generate mature B cells from hPSC-derived hematopoietic progenitors.

**Figure 6.**
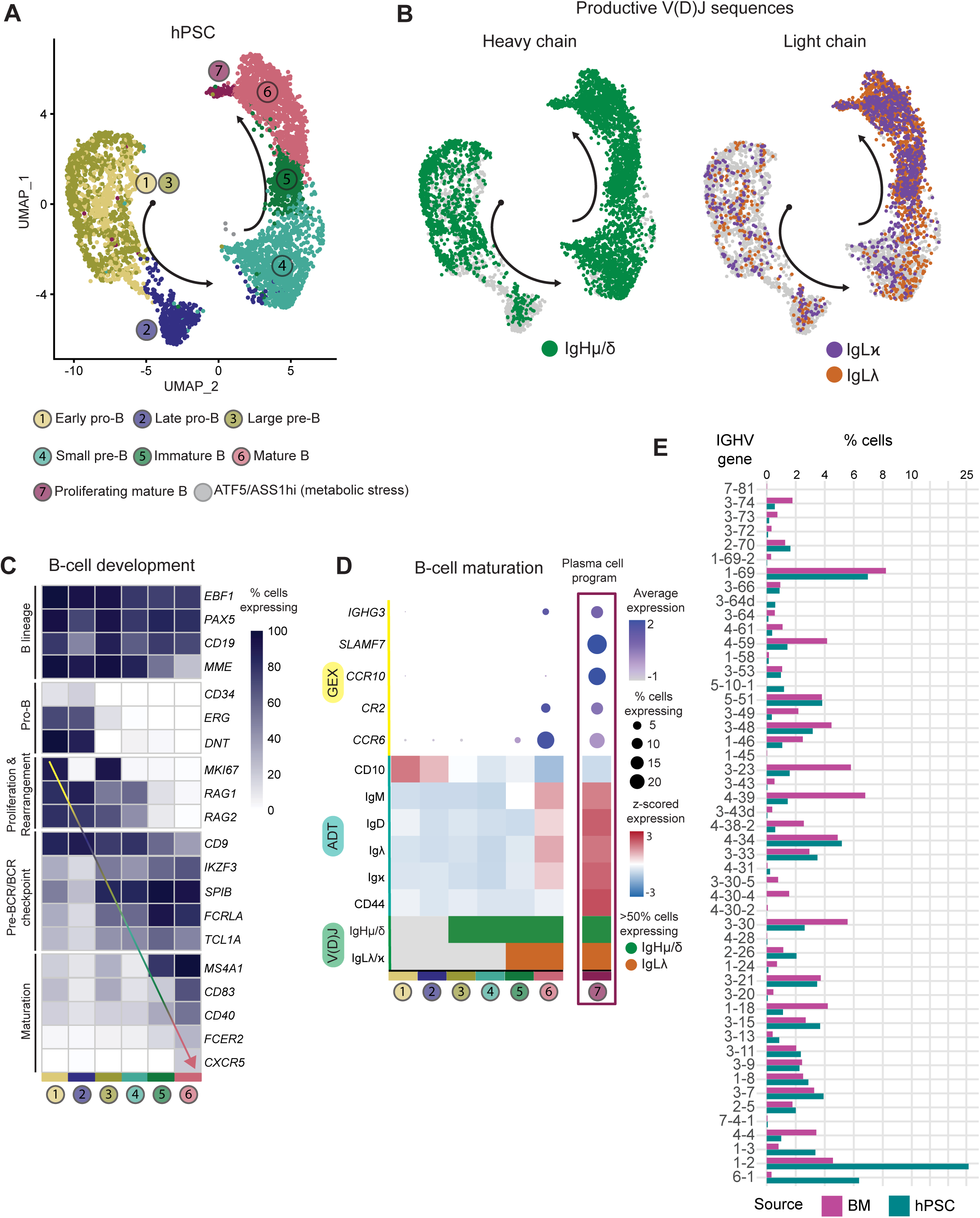
Multi-omic scRNA-seq analyses of adult bone marrow-derived and untreated hPSC-derived B lineage cells. (A) Harmony-integrated UMAP of total B lymphopoiesis from hPSC-derived B cells cultured without exogenous IL-7. Analyses were carried out on B cells (n=5,072 cells) isolated at day 35 of MS-5 co-culture. Arrows indicate the direction of B-cell development as determined by the 3-OMIC characterization (GEX, ADT, V(D)J). (B) Cells expressing productive immunoglobulin heavy chain (IgH) and light chain (IgL), kappa or lambda (IgL𝜅 or IgL𝜆), are indicated. hPSC-derived B cells show normal Ig V(D)J rearrangement (productive heavy and light chains), following canonical B-cell development. (C) Heatmap of the percentage of cells expressing selected genes (GEX) across B-cell developmental stages (Clusters 1 to 6). Genes are grouped to highlight key stages of B-cell development. The GEX heatmap shows that hPSC-derived B cells cultured in the absence of exogenous IL-7 follow a canonical B cell developmental pathway. (D) Dotplot showing percentage of cells expressing and the average log expression of genes associated with mature B cells (Clusters 1 to 6). Cluster 7 is highlighted (magenta outline) to indicate the most mature and proliferating cluster that contains cells expressing a plasma cell and IgG class-switched gene signature (*SLAMF7*, *CCR10*, *IGHG3*). The heatmap shows the z-scored expression levels of cell-surface markers (ADT) associated with B-cell maturation across the developmental stages (Clusters 1-7). (E) V(D)J repertoire analysis, comparing the immunoglobulin heavy chain variable gene (*IGHV*) usage between B cells from human bone marrow (BM) and untreated hPSCs. *V* genes are ordered based on their genomic location in reference to the *D* genes, from distal to proximal (obtained from IMGT).

To determine if the hPSC-derived mature B cells can further differentiate into functional plasma cells, we next cultured CD19^+^ cells isolated from day 35 cultures in microtiter wells under conditions that we have previously shown promote plasma cell differentiation of primary tonsillar naïve B cells.^48^ With this protocol the cells are maintained over a 10-day period in a sequence of four different culture conditions designed to mimic stage-specific signals that are received by B cells as they differentiate into plasma cells *in vivo*. By day 10 of culture, the hPSC-derived population contained CD27^+^CD38^+^ antibody-secreting cells, most of which had matured into CD27^+^CD38^+^CD138^+^ plasma cells (Figure 7A,B). Similar subpopulations were detected in cultures initiated with primary tonsillar naïve B cells. ELISPOT analyses revealed the presence of IgM, IgG, and IgA secreting cells, demonstrating that the hPSC-derived B cells had differentiated to plasma cells that had undergone antibody isotype switching (Figure 7C). ELISA analysis of the supernatants from the cultured cells showed the presence of IgM, IgG, and low amounts of IgA, confirming the presence of hPSC-derived plasma cells (Figure 7E). On a per B cell input, the levels of secretion were approximately 10-fold lower than those generated from the primary tonsil naïve B cell-derived plasma cells (Figure 7F). These findings clearly show that it is possible to generate functional antibody-secreting plasma cells from hPSC-derived B cells.

**Figure 7.**
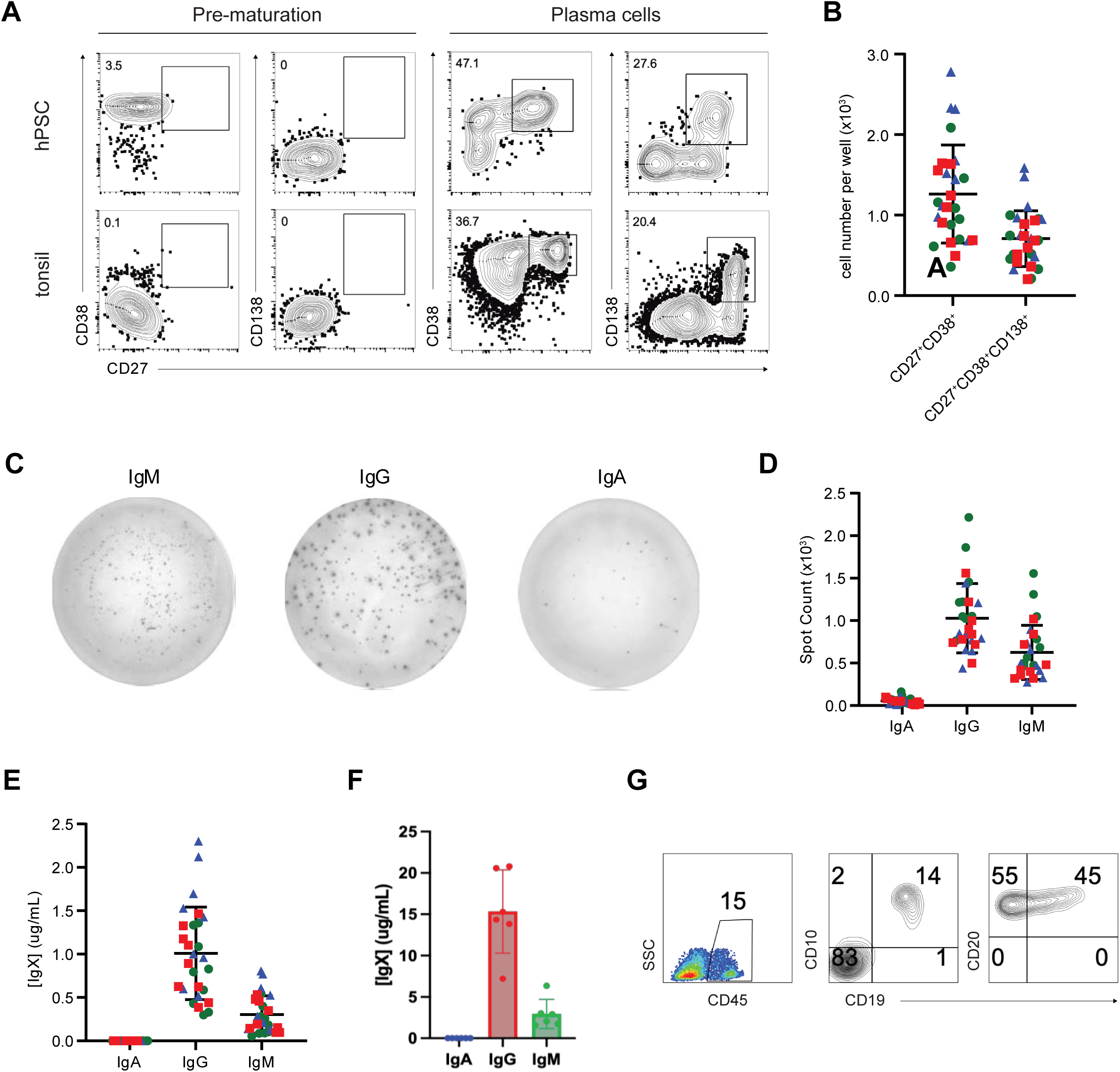
Generation of plasma cells from hPSC-derived B cells. (A) Representative flow cytometric analyses of expression of indicated markers on hPSC- and tonsil-derived cells prior to and following 10 days of culture (plasma). Tonsillar CD19^+^IgM^+^IgD^+^ naïve B cells were cultured directly after isolation, CD19^+^ hPSC-derived cells were isolated by FACS from populations cultured for 35 days on MS-5 stromal cells (B) Quantification of antibody-secreting cell numbers following 10 days of maturation culture. Each symbol represents a single well, different colors indicate different experiments (C) Representative isotype-specific ELISpot analyses of hPSC-derived plasma cells (D) Quantification of the number of antibody secreting cells (spots) for each isotype as determined by the ELISpot assay. As above, each symbol represents the antibody-secreting cell content of a single well, different colors indicate different experiments. (E) Quantification (by ELISA) of the amount of antibody detected in the supernatant of day 10 hPSC-derived plasma cell cultures. Shown are the amounts of the different isotypes detected (F) Quantification (by ELISA) of the amount of antibody detected in the supernatant of day 10 tonsil-derived plasma cell cultures. For the hPSC analyses, microtiter wells were seeded with 5x10^4^ CD19^+^ cells isolated from populations cultured for 35 days on MS-5 cultures. For all graphs, each symbol represents a single well and different colors indicate different experiments. Naïve tonsil-derived B cells were seeded at 10,000/well to avoid overgrowing the culture. Antibody concentrations were multiplied by a factor of 5 to “normalize” to 50,000 input cells used for the hPSC-derived B cells. (G) Flow cytometric analyses of CD45, CD19, CD10 and CD20 expression on MPP-derived populations cultured on MS-5 for 21 days.

### B cell potential distinguishes yolk sac primitive and MPP hematopoiesis

The above studies focused on the hPSC-derived definitive program and demonstrated that the HEC population has B-cell potential. In the next set of experiments, we mapped B cell potential across the human YS programs, to determine if, as in the mouse, these early developing populations also display the capacity to generate B lineage cells. For these studies we analyzed both the primitive and MPP programs using our previously described strategies.^22,27^ Definitive progenitors were included as controls.

In the initial studies, we evaluated the hPSC-derived yolk sac MPP program as we have previously shown that it has T cell potential and reasoned that it was the most likely population to display B cell potential. For these studies, the CD34^+^CD43^-^ fraction was isolated from day 6 EBs induced with BMP4 and activin A as previously described ^22^ (Fig. S7A). Following isolation, the cells were reaggregated and the aggregates maintained for 3 days in the B cell priming factors described above prior to culture on MS-5 stroma. The aggregates were then cultured on MS-5 for 21 days, with IL-7 added between days 7 and 21. As shown in Figure 7G, the MPPs did give rise CD19^+^CD10^+^CD20^+^ pre-B cells. The output was lower than that from the definitive progenitors [approximately 10^3^ cells per 5X10^4^ input], an observation consistent with the interpretation that they represent distinct populations.

Primitive hematopoiesis was specified through the staged inhibition of WNT signaling (Fig. S7B) a protocol that produces a transient wave of primitive progenitors that emerge between days 5 and 8 of differentiation and can be detected initially by the expression of CD34 and subsequently by the co-expression of CD34 and CD43 (Fig. S7C). To evaluate B cell potential, CD34^+^CD43^-^ progenitors were isolated daily (days 4-8) by FACS and cultured on MS-5 for 21 days. While CD45^+^ cells were generated from the day 4 and 5 progenitors, none of the populations from any of the time points gave rise to B cells (Fig. S7C). The CD45^+^ cells that developed were CD33^+^ myeloid cells (not shown). CD43^+^ hematopoietic populations isolated from days 8 culture also failed to generate B cells (not shown). By contrast, control definitive CD34^+^CD43^-^ progenitors isolated as early as day 6 of differentiation gave rise to CD10^+^CD19^+^ B cells (Fig. S7C). Taken together, these findings indicate that the hPSC-derived YS population does have B cell potential and that it is restricted to the MPP program.

## Discussion

The ability to generate specific blood cell types from hPSCs has provided a platform for mapping the early stages of embryonic hematopoietic development, for modeling diseases that target different progenitor populations and for producing functional cells for novel therapeutic applications. While most hematopoietic lineages can now be generated from hPSCs, efforts at differentiating B cells have met with limited success. Here we report a stepwise protocol for the generation of B cells from hPSCs and show that, as the cells differentiate, they transition through well-established stages of lineage development including pro-B and pre-B cell progenitors, naïve Ig-expressing B cells and finally Ig-secreting plasma cells. The production hPSC-derived plasma cells was one of the key goals of the study, as these cells represent the relevant target population for future cell-based therapies.^2^

To generate B lineage cells from hPSCs we focused our analyses on populations that we have previously shown contain T cell progenitors, specifically definitive HECs and YS MPPs.^22,24^ With this strategy, we gained the following new insights into the generation and characterization of hPSC-derived B cell progenitors that enabled us to produce B cells from multiple hPSC lines. First, we showed that B cell progenitors develop as early as day 8 of hPSC differentiation and are restricted to the CD34^+^CD184^-^CD73^-^ HEC fraction of this population, a pattern that recapitulates T lymphoid development. Second, we demonstrated that B cell potential of the CD34^+^CD184^-^CD73^-^ HEC fraction increases dramatically between day 8 and 13/14 of differentiation, suggesting that the definitive hematopoietic population is expanding over this time. Together with this increase in potential, the population appears to have matured as the day 13/14 progenitors generated B cells when plated directly on the MS-5 stroma, without the aggregation step required for the earlier cells. Our interpretation of these differences is that these HECs have already undergone the EHT in the EBs and as such, likely represent the earliest emerging hematopoietic progenitors. Finally, we showed that B cell potential was detected in the YS MPP population in addition to the definitive HEC population, suggesting that there are at least two developmentally distinct sources of human B cells. Whether these derivative B cells represent different B cell lineages, comparable to the B1 and B2 populations in the mouse is currently under investigation. Collectively, these findings provide the first developmental map of the human B lineage and in doing so, establish a foundation for further optimizing the generation of both the definitive and YS progenitor populations.

Generation of the B lineage cells from the hPSC-derived HECs appears to be largely driven by factors provided by the MS-5 stroma and those present in the fetal calf serum in the Myelocult media, as the cells differentiated and matured over a 35-day period without addition of exogenous cytokines. One pathway that may be involved is the IL-7 pathway, as studies have shown that MS-5 produces low amounts of IL-7 that can support human CB-derived B cell development.^19^ Given this and the findings that exogenous IL-7 can enhance human B lymphopoiesis in stromal cell cultures *in vitro*, we tested the effects of adding IL-7 during the specification and expansion of the pro/pre-B cell population (day 14 to 21) with the goal of improving B cell output from hPSCs. While we did observe an increase in the number of CD19^+^ pre-B cells generated in IL-7-treated cultures, the most striking effect of sustained IL-7 signaling was a complete block in lineage maturation, characterized by the emergence of B progenitors with gene expression patterns similar to pre-B cells, but unable to undergo productive IgH rearrangements. Surprisingly, these IgH-defective cells did generate productive IgL rearrangements, indicating that they could bypass the IgH developmental checkpoint in the absence of a functional pre-BCR. While this direct progression to IgL rearrangement without a productive heavy chain rarely occurs in normal B lymphopoiesis, it has been reported in certain conditions in mice^49^ and humans^50^. Notably, Shaw et al. showed that enforced expression of an activated RAS protein in J_H_ mutant pro-B cells unable to undergo IgH rearrangement promoted their differentiation and subsequent Ig Kappa rearrangement in a chimeric mouse model.^51^

The interplay between IL-7 signaling, proliferation, and Ig rearrangement is well documented in the mouse B cell lineage.^15^ Sustained signaling has been shown to prevent *Rag1* expression and inhibit IgL (Igκ) rearrangements in pre-B cells maintained *in vitro.*^52^ Our observation that IL-7 treatment in hPSC-derived B cells leads to both *RAG1* and *RAG2* expression (Figure 3) and functional IgL rearrangement in the absence of IgH suggests that the mechanisms in the human cells are likely different from those regulating Ig rearrangement in the mouse. Notably, the developmental block in the hPSC-derived population does not appear to be reversible, as cells subsequently cultured in the absence of IL-7 from days 21 to day 35 did not mature to express IgM and IgD (Figure 4A). Thus, it is possible that the sustained proliferative signal from the exogenous IL-7, through a developmental stage that normally exits the cell cycle to allow productive IgH rearrangements, is responsible for this abnormal differentiation program.

For hPSC-derived cell types to be useful for research and therapeutic applications, their development should follow a normal trajectory, and they must display appropriate functional properties. In this study, we used molecular and flow cytometric analyses and a multistep maturation/activation culture system to characterize, in detail, these aspects of the hPSC-derived human B cell lineage. Our transcriptomic analyses revealed striking similarities between the day 35 hPSC- and adult bone marrow-derived B-lymphopoiesis, demonstrating that the *in vitro* generated lineage transitioned through the expected normal developmental stages. The observation that staged inhibition of TGFβ signaling leads to an increase in the generation of IgM^+^IgD^+^ cells suggests that development/proliferation of the hPSC-derived B cell lineage, similar to the adult marrow lineage is antagonized by signaling through this pathway.^46^ Analyses of IgH recombination showed that hPSC-derived B cells exhibit diverse VDJ gene family usage indicating that they are not developmentally restricted in their ability to generate a broad BCR repertoire. These findings also demonstrate that this critical developmental stage is not dramatically impacted by the artificial stromal-based culture conditions. The diversity of *IGHV*, *IGHD*, and *IGHJ* genes detected in the B cells also shows that these cells are not the progeny of a few pre-B cell clones, further supporting the interpretation that the culture conditions support robust human B lymphopoiesis. Finally, the naïve B cells generated under these conditions responded to maturation/activation factors and differentiated into functional plasma cells capable of antibody secretion and class switching. Collectively, the outcome of these analyses demonstrates that the generation of hPSC-derived B cells follows the developmental path established for B lineage in the human adult bone marrow and generates cells that display functional properties of antibody-secreting plasma cells.

In summary, the findings outlined in this study advance our understanding of B cell development from hPSCs and show that though the manipulation of different signaling pathways, it is possible to generate different staged B cell populations including functional plasma cells. The ability to generate plasma cells represents an important step forward as it provides access to the population required for the development of novel cell-based therapies aimed at the long-term delivery of protein drugs or pathogen-specific antibodies to patients.

## Acknowledgment

We thank current and former members of the Keller lab for their advice and critical feedback on this work and the University Health Network/SickKids Flow Cytometry Facility for their assistance with FACS. The CHOPWT18 and RM TOM iPSC lines were obtained from Dr. Deborah French (CHOP, Philadelphia) and Dr. Elizabeth Ng (Murdoch Children’s Research Institute, Melbourne) respectively. The annotated adult bone marrow object was provided by Junkai Yang (Ghosn Lab). Next-generation sequencing services were provided by the Emory NPRC Genomics Core (RRID: SCR_026418), which is supported in part by NIH P51 OD011132. Sequencing data was acquired on an Illumina NovaSeq 6000, funded by NIH S10 OD026799. This work was supported by grants from the Canadian Institute of Health Research (CIHR) FDN159937 (G.K.), National Institutes of Health (NIH) R01AI099108 (D.B.), P30CA023074 (for the University of Arizona Flow Cytometry Shared Resource), and R01AI182276 (E.E.B.G.), and the Gates Foundation INV-051356 (D.B.), INV-071091 (D.B), INV-007910 (D.B.), INV-044064 (E.E.B.G.), NV-071644 (E.E.B.G.) and INV-050917 (G.K.)

## Author Contributions

G.M.K. and X.S. conceived the project, and X.S. designed and performed initial experiments and analyzed the data. X.S. and G.M.K wrote the manuscript. J.J.K., M.A. K.C.T., X.Z., V.M. and M.K. designed and performed experiments and J.J.K., X.Z. and M.K. analyzed data. K.K., A.F.N., A.K., and E.E.B.G. designed, performed, and analyzed multi-omics single-cell assays for the hPSC-derived and human bone marrow populations. B. Q. T. and D.B. performed the CITE-seq analyses of adult bone marrow, and C.A.F. and D.B. conducted and evaluated the maturation/activation studies on the hPSC-derived B cells.

## Methods

### Maintenance of hPSCs

The H1 hPSC line (Thomson et al., 1998) was maintained on irradiated mouse embryonic fibroblasts in hESC media containing DMEM/F12 (Cellgro) with non-essential amino acids (1x, Thermo Fisher Scientific), β-mercaptoethanol (55 µM, Thermo Fisher Scientific), penicillin/streptomycin (1%, Thermo Fisher Scientific), L-glutamine (2 mM, Thermo Fisher Scientific), KnockOut serum replacement (20%, Thermo Fisher Scientific) and 10 ng/mL FGF2 (R&D Systems). Media was changed daily for 5-6 days. hPSCs were passaged once at a ratio of 1:6-1:9 prior to differentiation. Cells were maintained in normoxic conditions (37°C, 5% CO_2_). The hPSC studies were subject to approval by the Stem Cell Oversight Committee (Canadian Institutes of Health Research). The CHOP18 ^47^ and RM TOM ^26^ hiPSC lines were maintained feeder free in E8 media.

### Hematopoietic differentiation of hPSCs

*Day 8 HECs and Yolk sac primitive progenitors*. For hematopoietic differentiation, hPSCs at 80-90% confluency were dissociated to generate embryoid bodies (EBs) as described previously ^22^. Briefly, the cells were treated with TrypLE (Thermo Fisher Scientific) to generate small clusters (< 5 cells per cluster) which were cultured at the equivalent of 500,00 cells/mL in 4 ml of StemPro-34 media (Gibco) in 60 mm Petri dishes (Falcon). The media was supplemented with base components including penicillin/streptomycin (1%, Thermo Fisher Scientific), L-glutamine (2 mM, Thermo Fisher Scientific), ascorbic acid (50 µg/mL, Millipore Sigma), transferrin (150 µg/mL, ROCHE), monothioglycerol (50 µg/mL, Millipore Sigma) (hereafter referred to as base media) along with ROCK inhibitor Y-27632 (10 µM) and BMP4 (1 ng/mL). The cultures were maintained on an orbital shaker (70 RPM) in a hypoxic environment (37°C, 5% CO_2_, 5% O_2_) for 18 hours to promote EB formation. Unless otherwise stated, the following differentiation steps were also carried out under hypoxic conditions.

Once formed, the EBs were harvested by centrifugation for 5 minutes and cultured in base StemPro-34 media supplemented with BMP4 (1 ng/mL) and FGF2 (5 ng/mL) for 24 hours. At this stage and for the duration of the differentiation protocol, the EBs were cultured under static conditions in polyhema (2-hydroxyethyl methacrylate) (Millipore Sigma)-coated tissue culture plates. On day 2, the EBs were harvested, washed and cultured further in base StemPro-34 media supplemented with BMP4 (10 ng/mL) and FGF2 (5 ng/mL). For induction of the primitive program, IWP2 (3 uM, TOCRIS) was added at this stage, while the combination of SB431542 (6 uM, TOCRIS) and CHIR99021 (3 uM, TOCRIS) were used to induce the definitive program. On day 3, the EBs were harvest, washed and cultured in base StemPro-34 media supplemented with VEGF (15 ng/mL), FGF2 (5 ng/mL), IL-6 (10 ng/mL) and IL-11(5 ng/mL) for 3 days at which point they were harvested and transferred to base StemPro-34 media supplemented with VEGF (15 ng/mL), FGF2 (5 ng/mL), IL-6 (10 ng/mL), IL-11(5 ng/mL), SCF (50 ng/mL), EPO (2 U/mL) and IGF-1 (25 ng/mL) for an additional 2 days. On day 8 of culture, the EBs were harvested, the cells dissociated, stained and subjected to FACS. The isolated populations were reaggregated in base StemPro-34 media supplemented with either the hematopoietic factors VEGF (5 ng/mL), TPO (30 ng/mL), IL-3 (30 ng/mL), IL-6 (10 ng/mL), IL-11 (5 ng/mL), SCF (50 ng/mL), EPO (2 U/mL), IGF-1 (25 ng/mL), BMP4 (10 ng/mL), FGF2 (5 ng/mL), FLT3L (20 ng/mL) or the priming factors TPO (30 ng/mL), IL-3 (30 ng/mL), FLT3L (20 ng/mL), G-CSF (20 ng/mL), SCF (50 ng/mL) in 96-well low-cluster plates (Corning) at 250,000 cells/mL for different periods of time. These aggregates were harvested and seeded onto MS-5 stromal cells as detailed below.

*Day 13 H1 HECs.* The following modifications were made for generation of the H1-derived day 13 HECs. EBs were generated in a modified base media consisting of 50:50 StemPro-34:IMDM supplemented with BMP-4 (4ng/ml) and the above components minus the ROCK inhibitor Y-27632. The cultures were maintained on an orbital shaker (70 RPM) in a hypoxic environment (37°C, 5% CO_2_, 5% O_2_) for 24 hours to promote EB formation. Cells were maintained under hypoxic conditions until day 9 of culture. On day 1, the EBs were harvested and transferred to the modified base media supplemented with BMP-4 (4ng/ml) and bFGF (5ng/ml) and cultured under static conditions in polyhema treated plates at an equivalent concentration of 1x10^5^ cells/ml. The definitive program was specified by the addition of SB431542 and CHIR99021 at day 1.75 of culture. On day 3.75, the EBs were harvested and dissociated to single cells with TrypLE (5 min, 37^0^C), washed two times, reaggregated in 6-well plates at a concentration of 2x10^5^ cells per ml in modified base media containing VEGF (10 ng/ml), bFGF (5ng/ml) and ATRA (1nM). The resulting aggregates were cultured under these conditions until day 6. At this time, ATRA was removed and the aggregates further cultured for 3 days in the supplemented base media with VEGF, FGF (as above) and IWP2 (3 uM). On day 9, IWP2 was removed and the aggregates were cultured in the supplemented media with VEGF, FGF (as above) under normoxic conditions. On day 11, the cultures were topped up with 1 ml/well of base media (minus VEGF/FGF). The cultures were harvested on day13 and processed as below.

*Day 13/14 HECs from iPSCs.* HECs were generated from the CHOP18 and RM TOM iPSCs using combined modifications of the above protocol and the protocol recently published by Ng. et al (2025). Mesoderm was induced in the EBs by treatment with CHIR99021 (3 µM) and rh FGF2 (30ng ml). 24 hours following initiation of mesoderm induction, the media was replaced with fresh media containing the TGFβ inhibitor SB431542 (6µM) along with CHIR99021 and rh FGF2. Following an additional 24 hours of culture, the EBs were harvested, the cells dissociated, then reaggregated and replated as above in media containing VEGF (50 ng ml-1), BMP4 (20 ng ml-1), rh FGF2 (20 ng ml -1) and RetAc (2 µM). Media was changed on day 4, 6, 9 and 11. Cytokine concentrations are modified as follows: the concentration of VEGF is lowered to 5 ng ml-1 at day 9 and to 1 ng ml-1 at day 11; the concentration of rhFGF2 is lowered to 5 ng ml-1 at day 9 and then maintained until day 14, BMP4 is lowered to 2ng ml-1 at day 9 and removed at day 11. RetAc was reduced to 500 nM on day 6 and to 200 nM on day 9 to day 14. In addition to the cytokines above, SR1 (20nM) and UM171 (35 nM) were added to the media at day 11. The cultures were harvested on 14 and processed as below.

*Yolk sac MPPs*. To induce the MPP (EMP/LMP) hematopoietic program, the day 1 EBs were collected by centrifugation at 40 RCF for 5 minutes and cultured further in StemPro-34 media supplemented with BMP4 (10 ng/mL), FGF2 (5 ng/mL) and Activin A (6 ng/mL) for 3 days. On day 4, the EBs were collected by centrifugation at 150 RCF for 5 minutes and cultured in StemPro-34 media supplemented with FGF2 (5 ng/mL), VEGF (15 ng/mL), IL-6 (10 ng/mL) and IL-11 (10 ng/mL) for 2 days. On day 6, the EBs were harvested, and the cells dissociated by Trypsin (Corning). The CD34^+^CD43^-^ fraction was isolated by MACS and/or FACS and the resulting cells were reaggregated in StemPro-34 media supplemented with TPO (30 ng/mL), IL-3 (30 ng/mL), FLT3L (20 ng/mL), G-CSF (20 ng/mL), SCF (50 ng/mL) and bFGF (5 ng/mL) in 96-well low-cluster plates (Corning) at 250,000 cells/mL for 3 days. These aggregates were used for B cell analyses. All recombinant factors are human and were purchased from R&D Systems.

### B cell differentiation of hPSC and cord blood progenitors

MS-5 stromal cells (kindly provided by Dr. K. Itoh at Kyoto University, Japan) were seeded into 6-well tissue culture plates (Corning) at a density of 10^5^-1.5x10^5^ cells per well in Myelocult H5100 medium (StemCell Technologies) supplemented with 1% penicillin/streptomycin and 2 mM L-Glutamine (supplemented H5100) 48 hours prior the initiation of B cell differentiation. For CB B cell differentiation, 10^3^ CD34^+^ CB cells (StemCell Technologies) were plated onto the MS-5 stroma in supplemented H5100 media and combinations of the following cytokines: TPO (50 ng/ml), SCF (100 ng/ml), IL-7 (20 ng/ml), IL-6 (50 ng/ml), FLT3L (10 ng/ml), IL-2 (10 ng/ml), GM-CSF (20 ng/ml), G-CSF (20 ng/ml) as outlined in the manuscript.

*D8 HECs and YS MPPs*: For hPSC definitive B cell differentiation, aggregates were generated by culture of day 8 HECs in 96 well low-cluster microtiter plates (Corning) (10^4^ cells per well) for 2 or 3 days in cytokine combinations indicated in the text. The aggregate contents of 1 microtiter well were plated directly onto 1 well of a 6-well plate seeded with MS-5 stroma and cultured in Myelocult H5100 media supplemented with 1% penicillin/streptomycin, 2 mM L-Glutamine, and with or without IL-7 (20 ng/mL) or TSLP (20 ng/mL) as described in the text. For MPP B cell differentiation, 6 CD34^+^ were isolated by FACS and the cells aggregated as above for 3 days. For these cells, aggregates from 3-5 microtiter wells were cultured in each well of a 6 well plate in the above conditions. IL-7 (20 ng/mL) was added at day 7 of B cell culture. For primitive and definitive B cell kinetics analyses, 3x10^4^ to 5x10^4^ CD34^+^ cells isolated (FACS) from day 4 to day 8 differentiated populations were cultured directly onto MS-5 stroma in the above conditions. IL-7 (20 ng/mL) was added at day 14 of B cell culture. The medium was changed weekly. For all populations, the developing B cells and stromal cells were harvested by trypsinization at day 21 of culture. The resulting population was filtered (40 μm strainer) to remove undissociated aggregates prior to further culture or analyses.

*D13/14 HECs:* CD34^+^CD184^-^CD73^-^CD43^-^ HECs isolated by FACS were seeded at different concentrations (5X10^2^-4.5X10^3^ cells per well) directly onto MS-5 stroma in 6 well plates containing supplemented Myelocult H5100 media with TPO, IL-3, G-CSF, FLT3L and SCF at the above concentrations. At day 3 of differentiation, the factors were removed and the cells cultured in media alone until day 14 of differentiation. In some experiments, IL-7 (20 ng/ml) was added between days 14 and 21. For extended cultures, SB (5.4 uM) was added from day 21 onward. The feeding schedule was as follows. On day 3 and day 7, the cells were cultured in supplemented Myelocult 5100 without additional growth factors. On day 14, the cells were cultured further in supplemented Myelocult 5100 with or without the addition of IL-7 as indicated. On day 21 the IL-7 was removed and, where indicated, SB (5.4 uM) was added to the cultures. The cells were maintained to day 35 in supplemented Myelocult 5100. The supernatant along with the non-adherent cells were aspirated from the wells before feeding on days 3, 7 and 14. On days 21, 28 and 35, the supernatant was collected and centrifuged at 336 x g and the pellet was mixed with fresh media and plated back into the wells. For days 24 and 31, fresh media was added directly to the wells at 2mL per well.

### Multi-omics single-cell assays

#### Cell enrichments, library preparations, and sequencing

hPSC-derived B cells (batch MBG7) were depleted of MS-5 cells using mouse CD29-PE antibody (Biolegend; Catalog #102208) and MojoSort™ Mouse anti-PE Nanobeads (Biolegend; Catalog #480080). MS-5-depleted cells were enriched for B cells using EasySep™ Human B Cell Enrichment Kit II Without CD43 Depletion (Stem Cell Technologies Catalog #17963). The hPSC-derived B cell batch MBG51 was enriched using CD19-positive selection (STEMCELL Technologies). Enriched B cells were encapsulated for single-cell RNA-sequencing using 10X Genomics Chromium Next GEM Single Cell 5’ Kit v2 kit (PN-1000263) to generate libraries for gene expression (GEX), B-cell receptor (BCR) V(D)J repertoire (PN-1000253), and cell-surface markers using antibody-derived tags (ADT) (PN-1000541) following the manufacturer’s protocol. The three libraries (GEX, V(D)J, ADT) were sequenced on an Illumina NovaSeq 6000 using an S4-200 flow cell (150k-200k mean reads per cell).

**For GEX**: FASTQ files for two hPSC-derived B-cells (MBG7, MBG51) and two adult bone marrow (ABM) donors (BM1, BM2) were processed using Cell Ranger v7.01 (10X Genomics) and mapped to the GRCh38-2020-A reference genome to generate gene expression count matrices. Matrices were further processed using R (v4.2.2) and Seurat (v 4.4.0). Count matrices were loaded with Read10X, and individual objects were made using CreateSeuratObject. For each object, PercentageFeatureSet was used to compute the percentage of mitochondrial gene expression (“^MT-” genes), and cells with >20% mitochondrial reads were excluded._Non-B cell populations were removed by excluding cells expressing markers of T cells (CD3E, CD4), myeloid cells (LYZ, MPO, ITGAM, ITGAX, CSF1R, MMP9), endothelial cells (CDH5), fibroblasts (CDH11), and mast cells (TPSAB1). The remaining cells were filtered for CD79A expression to enrich for B-cell lineage. Each dataset was normalized using NormalizeData and assigned a batch label in the metadata. The ABM reference data were obtained from our previous publication ^41^ in which we applied our “SuPERR” method of sequential biaxial gating to classify the various stages of B-cell development by integrating datasets from three OMICS (GEX, V(D)J, and ADT). Cell barcodes from high-confidence B-cell populations (clusters 1-6) annotated using SuPERR ^41^ were used to filter cells from the two adult BM datasets. Both filtered samples were merged to create a BM_total object, which served as a reference for mapping hPSC-derived B cells. **For V(D)J**: FASTQ files were processed using IMGT (International Immunogenetics Information System) (HighV-quest, v.3.6.1) and Cell Ranger (v.7.0.1, V(D)J reference – GRCh38-alts-ensembl-7.1.0). The IMGT output was processed using our VDJ analysis tool as previously described ^41^. The processed IMGT output was then merged by contig ID with the all_contig_annotations.csv file from the Cell Ranger output. The annotation files were loaded into R readr (v2.1.5) package. Contigs were filtered from the *high confidence* column of the Cell Ranger output, selecting those with at least two captured UMIs (Unique Molecular Identifiers). **For ADT**: To reduce background noise, we used the “dsb” (denoised and scaled by background v1.0) normalization method described by Mule et al. ^53^.

#### Dataset integration and batch correction

We integrated the samples using Harmony (v1.2.0). Data layers were joined using JoinLayers and log-normalized. Variable genes were identified using FindVariableFeatures(selection.method = “vst”, nfeatures = 3000). Immunoglobulin heavy and light chain (“^IG[HKL]”) and cell cycle genes from Seurat’s cc.genes were removed from the variable feature list. The data were then scaled using ScaleData. Principal components were computed using RunPCA, and batch effects were corrected using Harmony, grouped by the batch metadata. Harmony embeddings were used to calculate the nearest-neighbor graph, followed by clustering and UMAP coordinate generation using the functions FindNeighbors, FindClusters, and RunUMAP.

#### V(D)J annotation

A custom R function (annotate_vdj) was created to summarize the V(D)J annotations from the IMGT (International Immunogenetics Information System) and assign immunoglobulin (IG) categories as “productive” or “unproductive” for each cell barcode. If a cell contained both IG Kappa (IGK) and IG Lambda (IGL) contigs, then the IGL contig was retained, given that IGL rearrangement occurs after IGK. If a cell contained multiple contigs of either IGH or IGK/L, then the contig with the higher UMI count was retained—if the UMI counts were similar across the various contigs, that cell was filtered out as low-confidence or potential cell doublet. Each cell was classified into “productive”, “unproductive”, or “none” categories for its IGH, IGK, and IGL chains based on the v_domain_functionality field in the IMGT output. Annotated metadata was then added to the Seurat object using AddMetaData. After each sample was processed individually, they were combined into the Harmony-integrated Seurat object for visualization.

#### Code and Data availability

FASTQ, Cell Ranger output, and RDS files are available at NCBI GEO accession numbers GSE318326, GSE327194 (token for reviewers: *anahkocuvpaxjub*) and PRJNA1446811. The code used for data analysis and the associated figures are available at https://github.com/Ghosn-Lab

### Flow cytometry and fluorescence-activated cell sorting (FACS)

For surface marker analyses, single-cell suspensions were stained with fluorochrome-conjugated antibodies. Intracellular staining was performed with Cytofix/Cytoperm Fixation and Permeabilization Solution (BD), according to the manufacturer’s instructions. The following antibodies were used in this study: CD45-BV605, CD19-PECY7, CD10-APC, CD10-PE, CD33-FITC, CD33-APC, CD11B-APC, CD56-PE, CD56-APC, CD117-PE, CD34-PECY7, CD43-PE, CD43- FITC, CD73-PE, CD184-BV421, CD20-APC, CD20-PE, CD40-PE, CD123-PE, IgM-FITC, IgM-BV421, IgD-APC, Ig light chain lambda PE, Ig light chain lambda APC, Ig light chain kappa APC, CD38-PE, CD27-PE, CD45-APCCY7, CD4-PECy7, CD138-FITC, CD138-APC, sIgG-BB515, CD45-eF450. DAPI (Invitrogen) was used to stain dead cells. Stained cells were analyzed on a FACS LSR II (BD Biosciences) or LSRFortessa X-20 (BD Biosciences) analyzer. CD34^+^CD184^-^CD73^-^CD43^-^ HECs were isolated from either the day 8 or day 13/14 populations with a FACSAriaII (BD) cell sorter at the Sick Kids/UHN Flow Cytometry Facility. FlowJo version 10.8.1 (BD) was used for data analysis.

### Magnetic-activated cell sorting (MACS)

Single-cell suspensions were labeled with MACS Microbeads (Miltenyi Biotec) and the MACS separation was performed according to the manufacturer’s instructions. The following MACS Microbeads purchased from Miltenyi Biotec were used in this study: mouse cell depletion kit (130-104-694), human CD34 microbeads (130-056-702), human CD43 microbeads (130-091-333). Enriched cells were used for further culture or stained with antibodies for FACS or flow cytometric analysis.

### Real-time qPCR

Total RNA was isolated using the RNAqueous RNA Islolation Kit and treated with DNase I (Invitrogen). cDNA was prepared by reverse transcription using iScript Reverse Transcription Supermix (BioRad). qPCR was performed using the SsoAdvanced Universal SYBR Green Supermix (BioRad) on a Mastercycler Realplex^2^ (Eppendorf). Gene expression was normalized to TBP. Primers used in this study are listed in Supplementary Table 2.

### Maturation/activation

On day 0, 50,000 FACS-purified CD19^+^ B cells were seeded per well of 96-well U-bottom plates in IMDM supplemented with 10% fetal bovine serum (FBS, Peak), Antibiotic-Antimycotic (1ug/ml), ODN 2006 (50ng/ml), CD40L (50ng/ml), IL-2 (50ng/ml), IL-10 (50ng/ml), IL-15 (100ng/ml) and IL-21 (25 ng/ml) at 37C with 5% CO2. At Day 3, Rosiglitazone (CAS 122320-73-4) was added to a final concentration of 1uM. At Day 5, the media was replaced with IMDM supplemented with 10% FBS, Antibiotic-Antimycotic, CD40L (50ng/ml), IL-15 (100ng/ml), IL-21 (25ng/ml), IL-6 (50ng/ml), IFNg (10ng/ml), 1uM 2-NP (1uM), and NX-1607 (100 nM). At Day 7, the media was replaced with IMDM supplemented with 10% FBS, Antibiotic-Antimycotic, IL-2 (50ng/ml), IL-6 (50ng/ml), IL-10 (50ng/ml), IL-15 (100ng/ml), and IL-21 (25ng/ml). Cultures were harvested and analyzed on Day 10.

### Statistics

Data are presented as mean ± s.e.m. with all values from independent experiments. Statistical analyses were performed using one-way ANOVA tests with multiple comparison, two-way ANOVA tests and unpaired *t*-tests (two-tailed). GraphPad Prism version 11.0.0 was used for statistical analyses.

---

**Supplementary Figure 1.** B cell development from cord blood- and hPSC-derived progenitors. (A) Scheme of protocol used to generate B cells from CD34^+^ cord blood (CB) progenitors. (B) Representative flow cytometric analysis of CB-derived populations generated in the different cytokine combinations following 21 days of MS-5 co-culture. The CD19, CD10, CD33, CD56 profiles are from gated CD45^+^DAPI^-^ cells. 2G: GM-CSF, G-CSF. Cells cultured in the absence of factor represent the negative control. (C) Quantification of the frequency of CD19^+^ B cells, CD56^+^ NK cells and CD33^+^ myeloid cells generated in each of the indicated cytokine combinations *n* ≥ 3. (D) Left: Representative flow cytometric analyses of expression of indicated markers on day 8 CD34^+^ fractions [CD184^-^CD73^-^, HECs; CD184^+^CD73^lo^, arterial endothelial cells (AEC); CD184^-^CD73^+^, venous endothelial cells (VECs)] and on the populations generated from them following 21 days of culture on MS-5 stroma (right). The indicated populations were isolated by FACS, the cells aggregated with hematopoietic factors for 2 days and the aggregates cultured in the absence of factors (No factors). Right: Quantification of the number of CD19^+^ B cells generated from day 8 HECs, VECs, and AECs. *n* = 3. Error bars represent SEM. \**P* < 0.05, \*\**P* < 0.01, \*\*\**P* < 0.001, \*\*\*\**P* < 0.0001, ^##^*P* < 0.01, ^###^*P* < 0.001, ^####^*P* < 0.0001 by one way ANOVA analyses with Tukey’s multiple comparisons.

**Supplementary Figure 2.** Effect of IL-7 on B cell development from hPSC-derived progenitors. (A) Scheme of the protocol used to test the effect of IL-7 or TSLP on the generation of CD19^+^ cells from day 8 HECs. The factors were added during the indicated time intervals. (B) Quantification of the frequency of CD19^+^ B cells and of the total number and frequency of CD56^+^ NK cells generated from day 8 hPSC-derived HEC cultured for 21 days under the indicated conditions. Error bars represent SEM. \**P* < 0.05, \*\**P* < 0.01, \*\*\**P* < 0.001, \*\*\*\**P* < 0.0001, ^#^*P* < 0.05 by one way ANOVA analyses with Tukey’s multiple comparisons. (C) Scheme of the protocol used to generate B cells from day 13 HECs. Stages at which cytokines/pathway inhibitors are add are indicated

**Supplementary Figure 3.** Differential expression of pre-BCR program genes in adult BM-derived and IL-7 treated hPSC-derived B lineage cells. Split violin plot comparing the log-normalized expression of pre-BCR program genes in the large pre-B stage (cluster 3). Select genes involved in the pre-BCR checkpoint. These genes are upregulated during progression from pre-BCR to maturation, as shown for the human BM sample. BM=215 cells; IL-7 treated hPSC=1,486 cells.

**Supplementary Figure 4.** Regulation of B cell development in the MS-5 cultures. (A) Representative flow cytometric analyses of expression of indicated markers on day 21 hPSC-derived populations cultured in the indicated concentrations of IL-7. IL-7 was added to cultures between days 14 and 21. (B) Quantification of the total number of cells, the frequency of CD19^+^IgM^+^ cells and the frequency of CD19^+^CD10^+^ cells generated on days 21, 28 and 35 of culture in the presence of the indicated amounts of IL-7. (C) Violin plots showing surface protein expression levels of IgM and IgD across Seurat-defined clusters. (D) Violin plots showing expression of *TGFBR1*, *TGFBR2* and *TGFβ1* in day 21 hPSC-and bone marrow-derived B cell populations (E) Quantification of the frequency of IgM^+^IgD^-^ and IgM^+^IgD^+^ B cells and of the total number of cells generated on days 28 and 35 in the absence and presence of SB. SB was added to the cultures at day 21. Error bars represent SEM. \**P* < 0.05, \*\*\*\**P* < 0.0001 by two-way ANOVA analyses with Tukey’s multiple comparisons. N=4

**Supplementary Figure 5.** Generation of B cells from different iPSC lines. (A) Representative flow cytometric analyses of expression of indicated markers on B lineage cells generated from CHOP18 iPSC-derived HECs cultured for 21, 28 and 35 days. (B) Quantification of the number of total, CD19^+^CD45^+^ and CD19^+^IgM^+^ cells and of the frequency of CD19^+^CD45^+^ and CD19^+^IgM^+^ cells on the different days. Numbers shown are per well of a 6-well plated seeded with 1500 day 14 CD34^+^CD184^-^CD73^-^ HECs. Different symbols represent B cells generated from three independent HEC populations, the same symbols with different colors indicate B cells generated from the same HEC population. (C) Representative flow cytometric analyses of expression of indicated markers on B lineage cells generated from RM TOM iPSC-derived HECs cultured for 21, 28 and 35 days. (D) Quantification of the number of total, CD19^+^CD45^+^ and CD19^+^IgM^+^ cells and of the frequency of CD19^+^CD45^+^ and CD19^+^IgM^+^ cells on the different days. Numbers shown are per well of a 6-well plated seeded with 1500 day 14 CD34^+^CD184^-^CD73^-^ HECs. Different symbols represent B cells generated from three independent HEC populations, the same symbols with different colors indicate B cells generated from the same HEC population.

**Supplementary Figure 6.** Restored large pre-B phenotype and Ig V(D)J gene usage in untreated hPSC-derived B cells. (A) Split violin plot comparing the log-normalized expression of pre-BCR program genes in BM-derived (n=215 cells) and untreated hPSC-derived (n=1,034 cells) large pre-B cells (Cluster 3). (B) V(D)J repertoire analysis, comparing the immunoglobulin heavy chain diversity (*IGHD*) and immunoglobulin heavy chain joining gene (*IGHJ*) family usage between B cells from human bone marrow (BM) and untreated hPSCs. (C) Feature plots highlighting key stages of B-cell development in the untreated hPSCs-derived day 35 population (n=5,072 cells) including early pro-B (*MKI67*, *CD34*, *RAG1*), late pro-B (*CD34*, *DNTT*), large pre-B (*MKI67*, *RAG1*), small pre-B (*RAG1*, *IGF2*), immature B (*CD40*, *MS4A1*) and mature B (*CD40*, *MS4A1*, *CXCR5*, *MME*-ve).

**Supplementary Figure 7.** B cell potential of the primitive and MPP hematopoietic programs. (A) Schematic of the protocol used to induce MPP hematopoietic progenitors from hPSCs and the manipulations used to assay B cells from them. For these analyses, 10^4^ FACS isolated day 6 CD34^+^CD43^-^ MPP HECs were reaggregated in the presence of the priming cytokines with bFGF for 3 days. These aggregates were then replated onto MS-5 stroma and cultured in the indicated conditions. (B) Schematic of protocol used for generation of the primitive and definitive hematopoietic programs from hPSCs. IWP-2 and SB431542+CHIR99021 were used for primitive and definitive mesoderm induction respectively. (C) Representative flow cytometric profiles showing the emergence of primitive CD34^+^ and CD43^+^ and definitive CD34^+^ populations between days 3 and day 8 of differentiation. CD34^+^CD43^-^ HECs were isolated by FACS from days 4 to 8 of culture and assayed for B cell potential. (D) Representative flow cytometric analysis showing the development of CD45^+^ and CD19^+^CD10^+^ B cells populations generated from primitive (left panel) and definitive (right panel) FACS isolated CD34^+^CD43^-^ progenitors. CD19/ CD10 profiles are from gated CD45^+^ cells (top).

