## Supplementary Figures for "Modeling human B cell development with pluripotent stem cells"

### Supplementary Figure 1

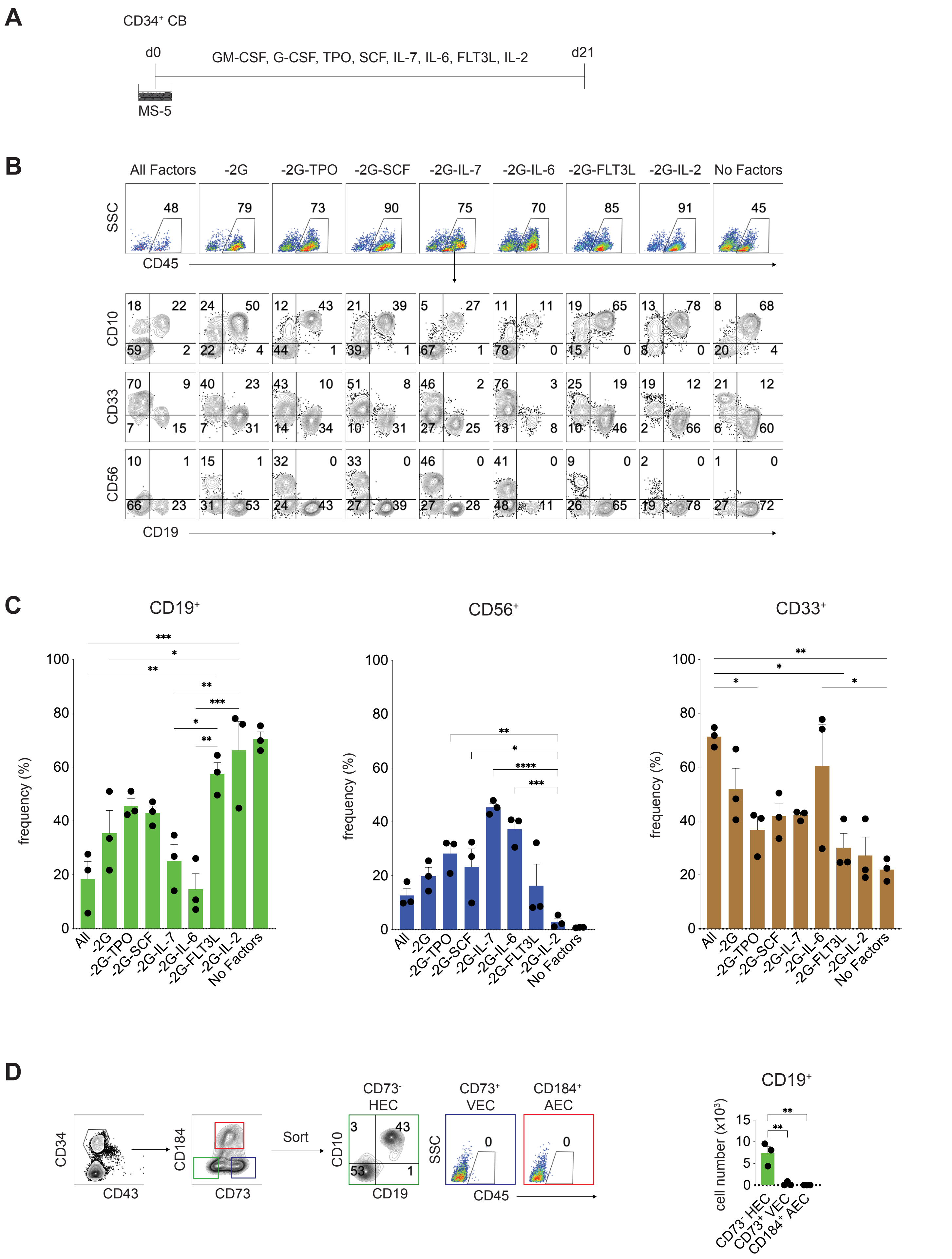

A

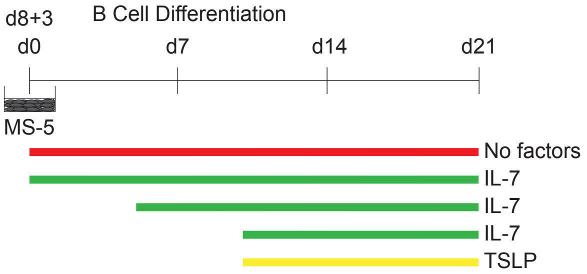

d8+3
B Cell Differentiation
d0
d7
d14
d21
MS-5
No factors
IL-7
IL-7
IL-7
TSLP


B

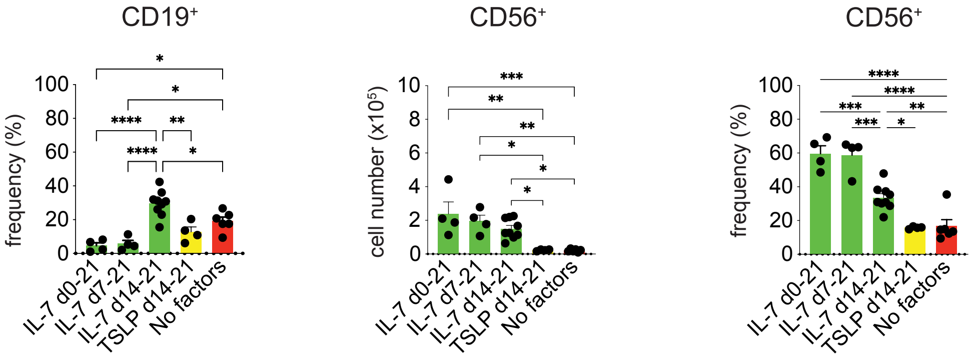

CD19+
frequency (%)
IL-7 d0-21
IL-7 d7-21
IL-7 d14-21
TSLP d14-21
No factors
CD56+
cell number (x10^5)
CD56+
frequency (%)


C

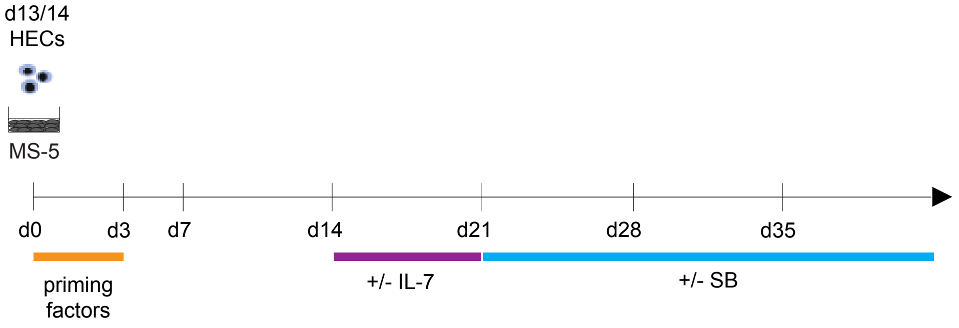

d13/14
HECs
MS-5
d0
d3
d7
d14
d21
d28
d35
priming factors
+/- IL-7
+/- SB

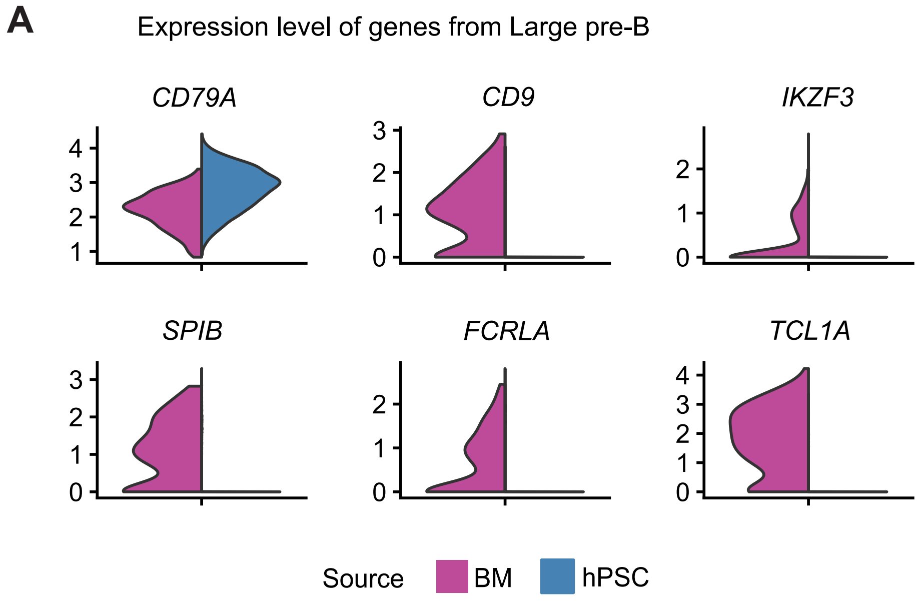
A
Expression level of genes from Large pre-B
CD79A
CD9
IKZF3
SPIB
FCRLA
TCL1A
Source
BM
hPSC

### Supplementary Figure 4

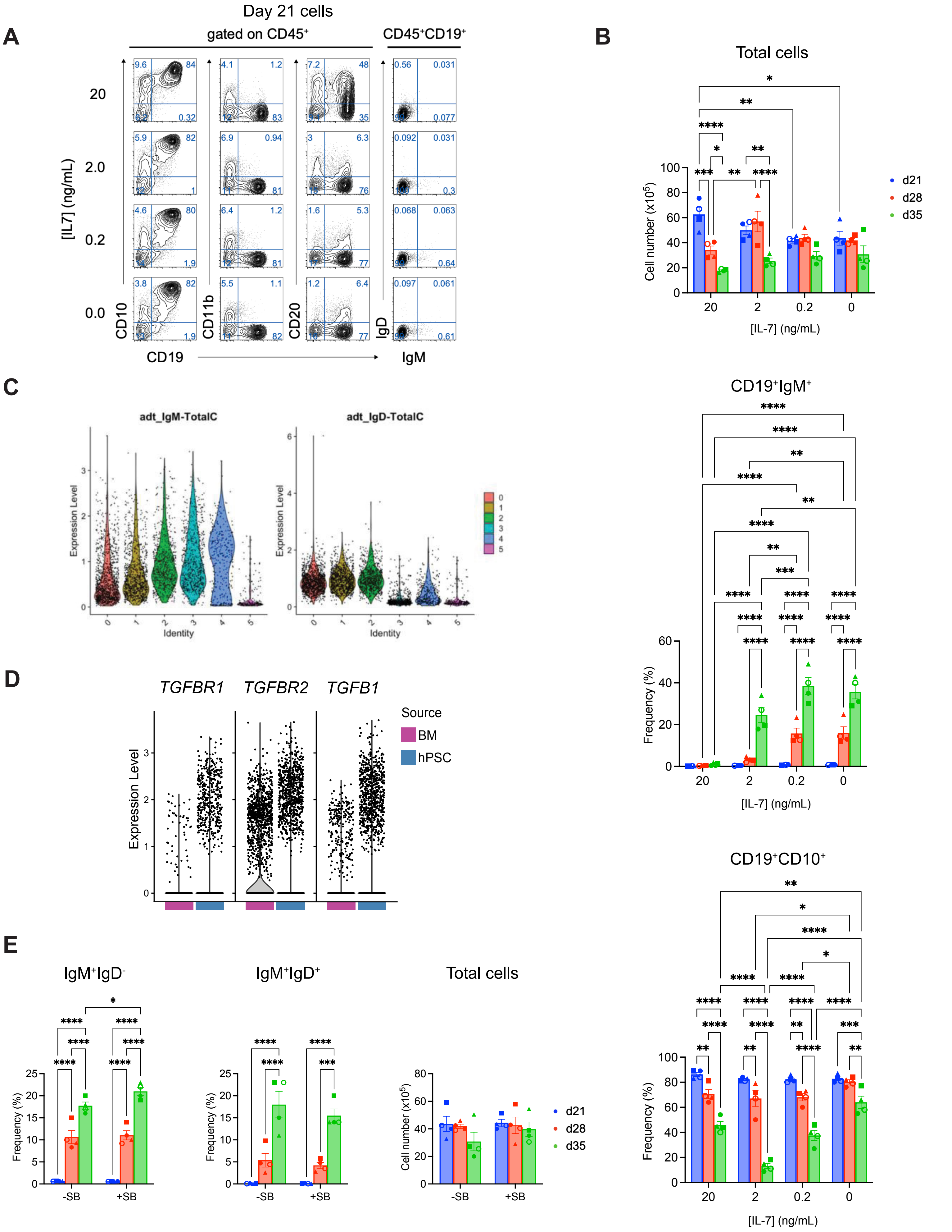

### Supplementary Figure 5

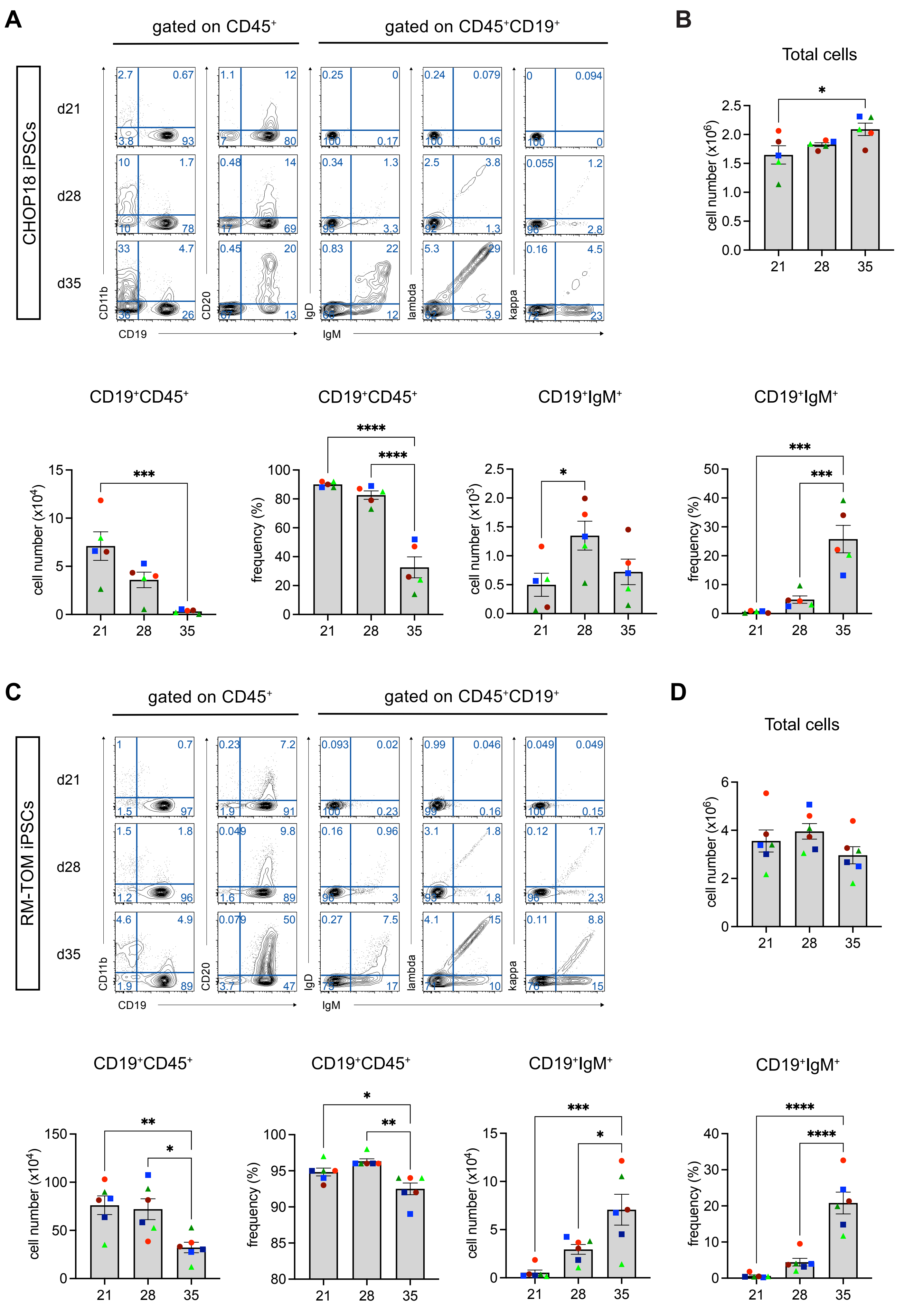

#### Supplementary Figure 6

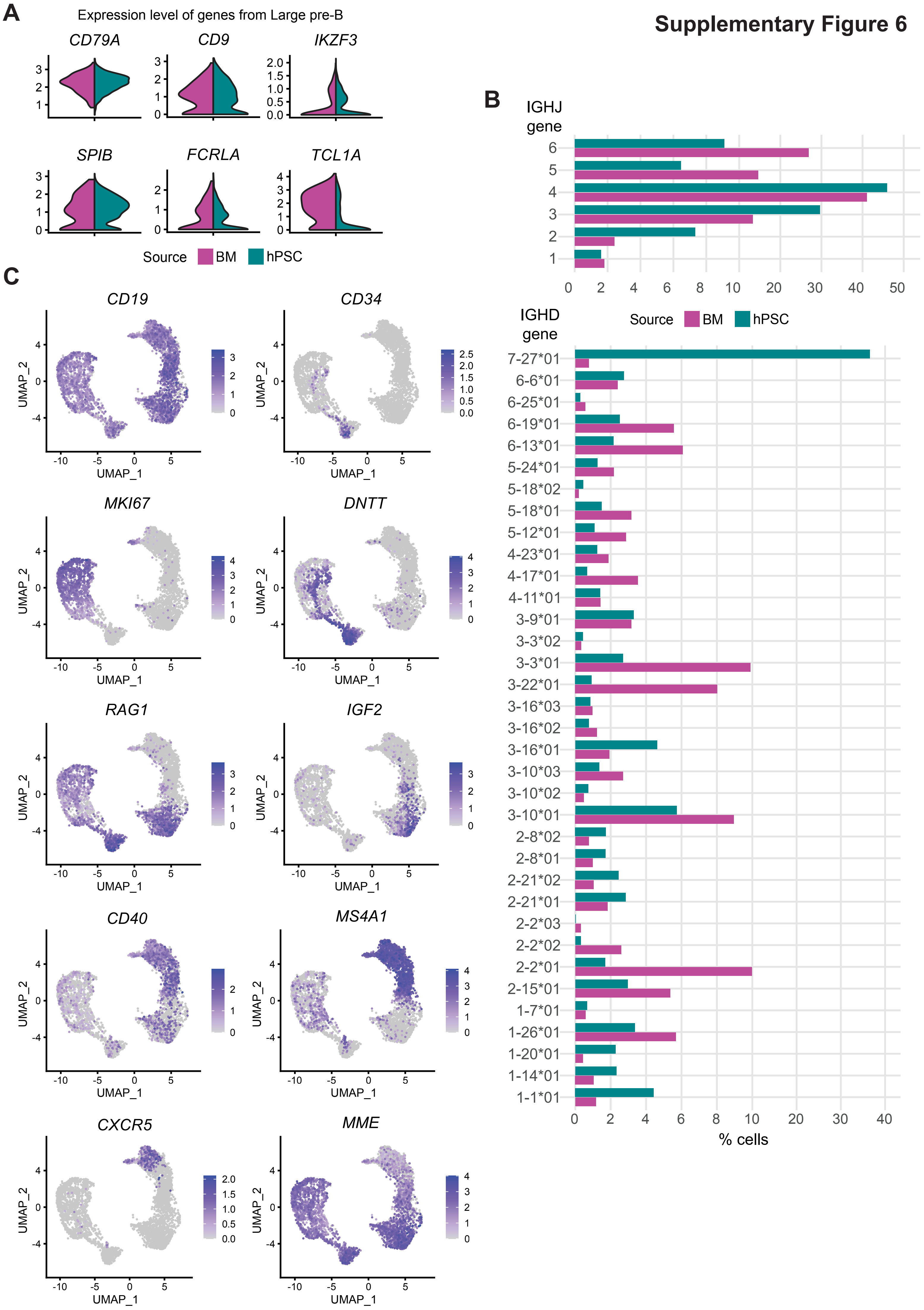

### Supplementary Figure 7

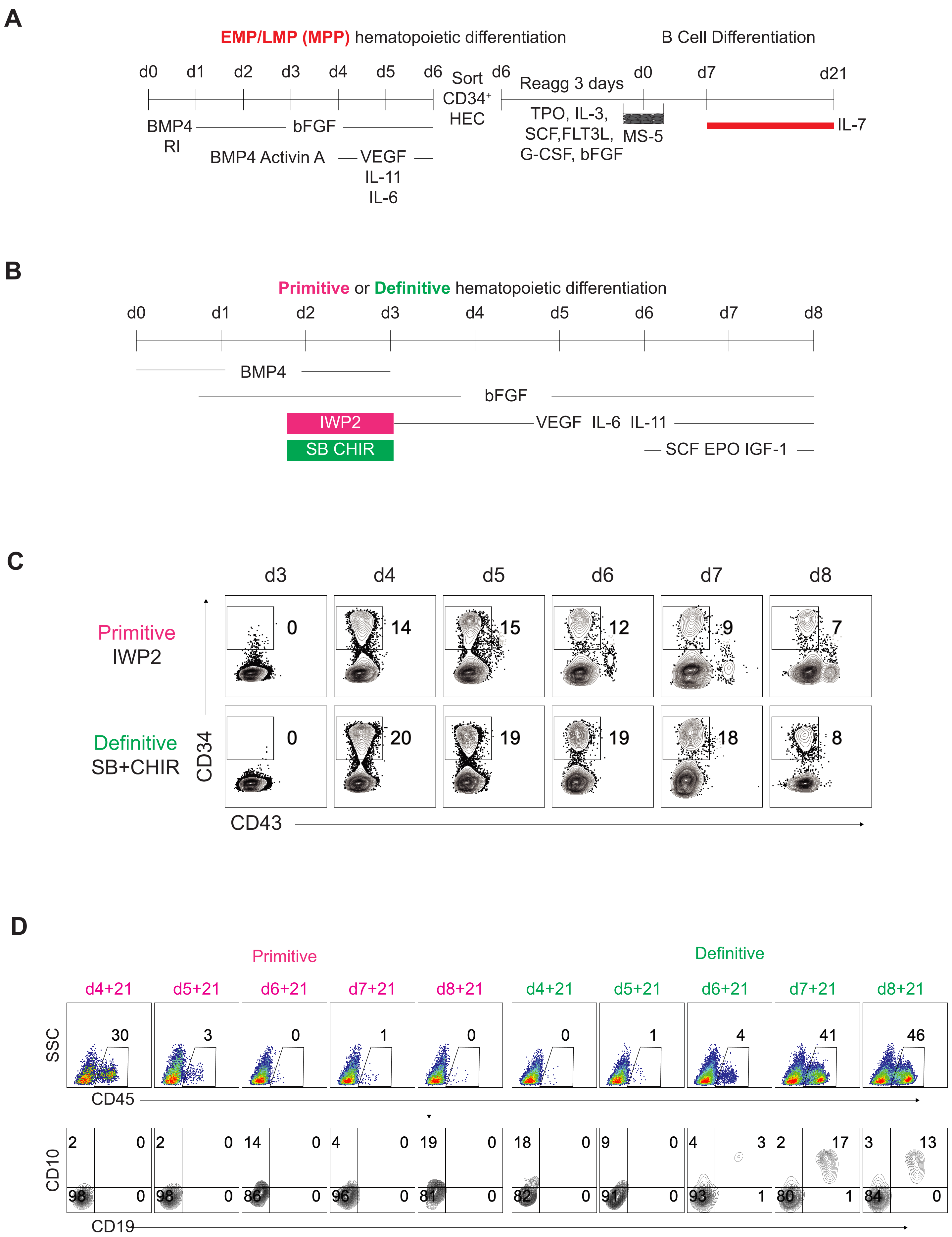
